# Homecage-Like Monitoring Reveals Domain-Specific Rescue of Central *SPTAN1* R1098Q Phenotypes by Wild-Type Partner Co-Housing

**DOI:** 10.64898/2026.09.25.754160

**Authors:** Patrycja Ziuzia, Michał Ślęzak, Arkadiusz Miążek

**Affiliations:** Research Group Biology of Astrocytes, P4Health Center, Lukasiewicz Research Network - PORT Polish Center for Technology Development, Wroclaw, Poland; Department of Biochemistry and Molecular Biology, Wroclaw University of Environmental and Life Sciences, Wroclaw, Poland

**Keywords:** *SPTAN1 R1098Q*, Computational Ethology, SLEAP, deepOF, SAM2, Home-Cage Monitoring, Social Buffering, Nest Building, 3R Principles

## Abstract

Neurodegenerative disorders manifest with markedly different phenotypes, with motor- and memory impairments dominating both clinical and preclinical phenotyping efforts. However, neurodegeneration in unlikely to be confined to these two functional domains alone, and its broader behavioral consequences, including social alterations remain underexplored. Here, we present a non-invasive, high-content home-cage phenotyping framework to investigate social and exploratory dysfunction in *SPTAN1* R1098Q mice, a model of early infantile epileptic encephalopathy (EIEE5). Our pipeline integrates three open-source AI engines: SLEAP.ai for markerless multi-animal pose estimation, DeepOF for automated classification of autonomous and social behaviors, and SAM2 for segmentation and quantification of nest-building material usage. Long-term monitoring revealed that *SPTAN1* R1098Q mutants exhibit impaired spatial habituation in mutant-mutant (MUT - MUT) pairs. Mutants also displayed persistent, unreciprocated trailing (following) of cage mates and a pronounced reduction of nest-building behavior alongside. Notably, co-housing with a wild-type littermate (MUT - WT) attenuated mutant hyperactivity and restored more typical rest pattern, suggesting that social context can partially buffer these behavioral abnormalities. By revealing social and exploratory alterations that expand conventional phenotyping, this computational ethology pipeline advances 3R principles while revealing that intrinsic executive state-switching failures in neurodegenerative models can be functionally buffered by social dynamics.

## Introduction

Spectrins are highly conserved structural proteins known to be primarily involved in the formation of the erythrocyte cortical membrane skeleton (Derbala et al., 2018). Among seven genes encoding mammalian spectrins, two encode α chains (α-I and α-II, encoded by the *SPTA1* and *SPTAN1* genes, respectively) and five encode β chains (β-I to β-V, encoded by *SPTB*, *SPTBN1*, *SPTBN2*, *SPTBN4*, and *SPTBN5*) (Liu & Rasband, 2019). While β-spectrins are ubiquitously expressed in the mammalian nervous system (Lorenzo et al., 2023), the two α-spectrin isoforms show markedly different distributions: α-I spectrin expression is restricted to erythroid cells (Wang et al., 2018), whereas α-II spectrin is expressed ubiquitously, with the highest levels found in the central nervous and cardiovascular systems (Skrzymowska et al., 2021). Within neurons and glial cells, spectrins contribute to building the membrane-associated periodic skeleton (MPS) — a ∼190 nm periodic structure first identified in neurons using super-resolution microscopy (Xu et al., 2013), later demonstrated to be found in oligodendrocytes (D’Este et al., 2016).

Spectrins act as actin-binding proteins, anchoring the cytoskeleton to cellular membranes directly or, in partnership with ankyrins — particularly Ankyrin-G and Ankyrin-B — indirectly (Huang & Rasband, 2018; Lorenzo, 2020). They are also known to play a key role in axonal transport, facilitating the movement of lipid-and protein-containing vesicles and organizing membrane-associated signalling complexes to maintain neuronal structure, polarity, and function (Liu & Rasband, 2019; Morrow & Stankewich, 2021). Given this central involvement in neuronal architecture and trafficking, specific mutations in spectrin genes lead to severe neurological dysfunction, affecting both, the central and peripheral nervous systems (Miazek et al., 2021; Zalas et al., 2023) — including early infantile epileptic encephalopathy (*SPTAN1*; Liu & Rasband, 2019) and Spinocerebellar Ataxia Type 5 (*SPTBN2*; Lise et al., 2012; Lorenzo, 2023).

Since spectrins’ primary aminoacid sequences are highly conserved across evolution, mice offer relevant animal models for studying inherited human spectrinopathies (Baines, 2009, Saitsu et al., 2010, Zalas et al., 2023). To date, however, characterization of these models has been dominated by motor coordination and balance deficits (Table 1, see Discussion), consistent with the broader tendency in neurodegeneration research to focus on motor and memory impairments at the expense of other, possibly affected domains. Reported examples of less-explored phenotypes include anxiety-like behavior, altered exploratory activity, and impaired nest-building in the mouse model carrying R1098Q point mutation (Hobbib et al., 2025), and reduced social behavior in a β-II spectrin (*Sptbn1*) heterozygous model (Cousin et al., 2021). Critically, these domains have been assessed using short-term motor or cognitive tests which often fail to capture complex, socially embedded behaviors that emerged only over prolonged conspecific encounters (Wang et al., 2016; Yang et al., 2012). Homecage-like monitoring offers measurements of complex behavior in more naturalistic environment, while minimizing confounding factors inherent in artificial laboratory environments, such as forced handling and social isolation (Puścian et al., 2014; Kizielewicz et al. 2026). Long-term recording in more ethological conditions enables to capture individual variability, spontaneous social dynamics, and subtle traits, often missed by short-term, traditional assays (Mitchell et al., 2020; Rusu et al., 2023; Liu et al., 2024).

**Table 1:**
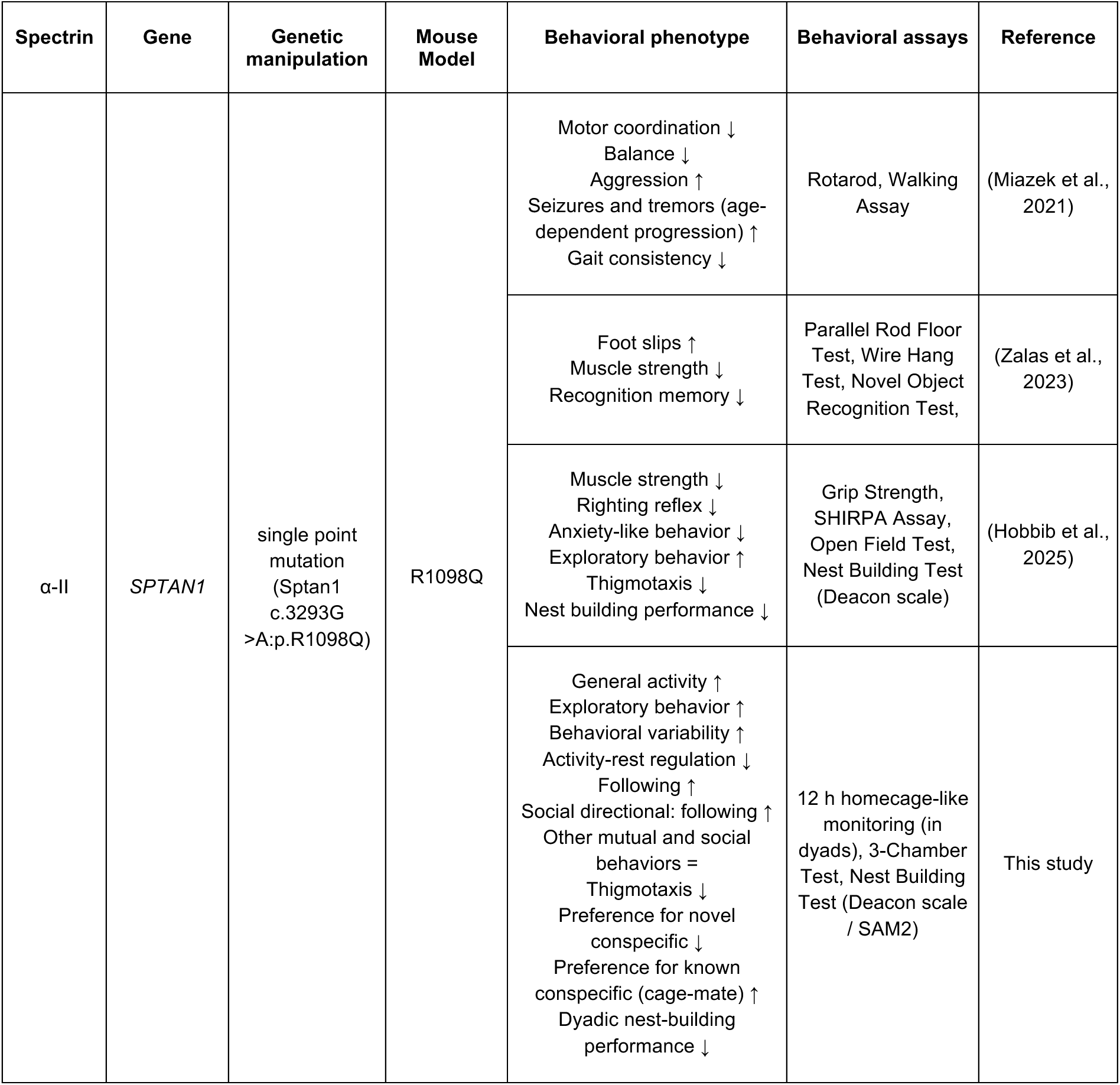
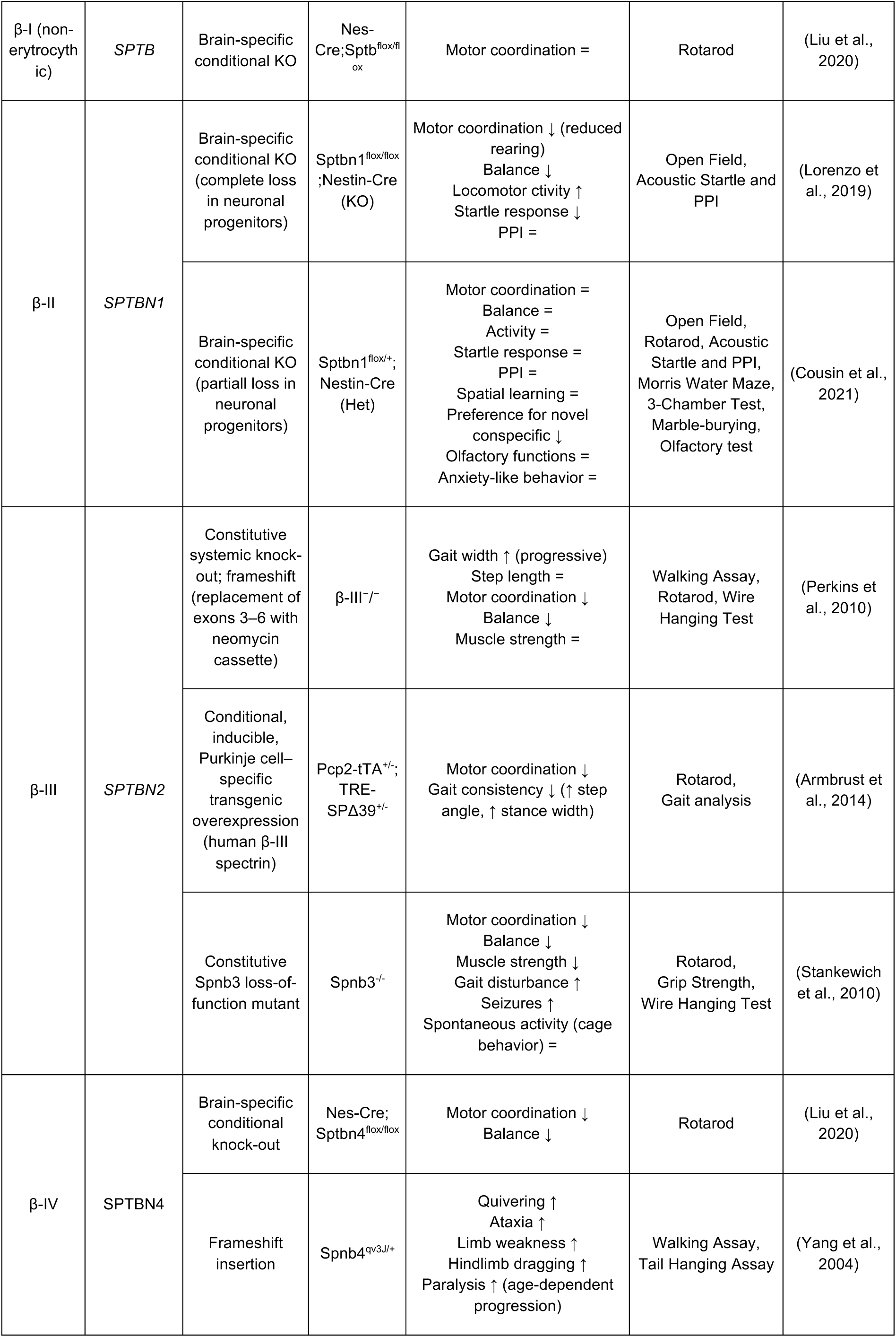
Summary of reported phenotypic characteristic of mouse models spectrinopathies. For behavioral assessment: ↑ increased; ↓ decreased; = unchanged; Behavioral phenotypes are reported relative to appropriate control littermates within cited reference.

Apart from autonomous and social behavior monitoring, nest-building activity is one of the most ethologically robust and evolutionarily conserved behaviors in rodents, known to be affected in R1098Q mice. The quantification of nest building activity has been based on endpoint scoring systems, but not the temporal dynamics of the nest construction (Deacon, 2006). Efforts to move beyond this limitation have primarily relied on specialized 3D depth-sensing hardware to extract structural nest parameters such as volume, wall height, and radius (Toyoda et al., 2015), revealing strain-specific differences in nest architecture and building speed. However, such approaches have not been used to monitor nest building with preservation of the social contexts, e.g. co-housed dyads.

This study was designed to characterize spontaneous behavioral dynamics and nest building of R1098Q mice (MUT) in home-cage-like environment. We further exploited the AI-based analytical workflow to assess behavioral profile of MUT and WT dyads. Classical behavioral paradigms were used to validate the core findings obtained from continuous observations of home-cage behavior (Shemesh & Chen, 2023). We discovered mutual effects on behavioral profile and nest building activity elicited by WT and mutant animals. This study expands the scope of home-cage monitoring and provokes further research on the normalization of neurodegenerative phenotype originating from mutation through social interactions with unaffected partner.

## Materials and Methods

### Animals

#### 3Rs

In accordance with the 3Rs principle, animals were drawn from existing breeding colonies rather than a dedicated breeding scheme. This strategy prevented excess animal production while ensuring adequate group sizes (Reduction). Standard, often overloaded behavioral test battery was replaced with continuous observation in a home-cage-like experimental arena. At least three environmental enrichment items were provided per cage, rotated according to subjects’ preference, and testing included both sexes to minimize overall animal use. Protocols were designed to minimize stress and the number of animals used (Refinement).

### Housing

All animal experiments were conducted in accordance with the European Union directive on the protection of animals used for scientific purposes (2010/63/EU) and approved by the Local Ethics Committee of the Institute of Immunology and Experimental Therapy PAS in Wroclaw, Poland (app. Nb. 044/2024, 17.07.2024). Experiments were carried out at the Animal Facility of Wroclaw University of Environmental and Life Sciences, Poland, under a 12:12 light cycle (lights off 7 AM–7 PM), controlled temperature (22 ± 2°C) and humidity (55% ± 10%). Mice were group-housed (2–4 per cage) in conventional cages (1284L Static Cage, Techniplast, Italy), 36,5 x 20,7 x 14 cm (L×W×H, 755,55 cm² floor area) with ad libitum access to water and food.

Behavioral tests were performed on both males and females on a C57BL/6J background, originating from breeding of Sptan1 c.3293G>A (p.R1098Q) mice (Miazek et al. 2021). To ensure phenotype consistency, mice were 6–10 months old at the start of experiments. This range corresponds to the fully established neurological symptoms of ataxia, reaching full development and stability between 6 and 9 months of mice age (Miazek et al. 2021, Zalas et al. 2023). Both males and females were included, however sex was not analyzed as a statistical factor, as previous studies shown insignificant gender differences (Miazek et al. 2021) The experimenter performing manual proofreading of tracked data was blind to genotype.

### Behavioral tests

#### Monitoring of spontaneous behavior in home-cage setting

Behavioral phenotyping was conducted in custom-built arenas (material: dibond) designed to resemble a home-cage-like environment, as previously described (Dalbeyler et al. 2025, bioRxiv), with minor modifications. Arenas measured 25 × 25 × 60 cm (L×W×H, 625 cm² floor area) and were separated by thick black curtains to prevent light and sound interference. Each arena was equipped with a food and water dispenser, bedding material, and three enrichment items (sunflower seeds hidden in bedding, a wooden block, two cotton-pressed nesting pads). Mice were introduced to the arena at the onset of the active phase (ZT0) and allowed to freely explore for 12 h (until ZT12) under infrared lighting.

Each arena was illuminated with infrared light (940 nm) via two LED plates (3 W, DC12V, 300 mA) mounted 80 cm above the floor. Video was recorded with Basler daA1920-160um cameras and Fujinon 6 mm f1.8 C125-0618-5M lenses (top-down view), controlled by a custom Python script allowing simultaneous recording of two arenas. Videos were saved as .avi, 648×600 px, 25 fps, grayscale.

Animals were always paired with their existing cage-mate, regardless of litter origin but co-housed prior to the procedures. One animal per pair was tail-marked for visual discrimination throughout the 12-h recording. Dyads were allocated as WT - WT (n = 5 boxes), WT -MUT (n = 10 boxes), and MUT - MUT (n = 4 boxes), yielding four individual-animal groups: WT (with WT, n = 10), WT (with MUT, n = 10), MUT (with WT, n = 10), MUT (with MUT, n = 8).

### Pose estimation and behavioral quantification

#### a. SLEAP.ai

Pose estimation was performed using SLEAP v1.6.1 (sleap-io v0.6.4, sleap-nn v0.1.0), Python 3.13.12, macOS 15.6.1 (Apple Silicon M4, 16 GB RAM). Two sequential models were trained following the top-down multi-animal pipeline (centroid model followed by centered-instance model), on 293 manually labeled frames from 7 randomly picked videos. Pixel-to-centimeter conversion (0.090232 cm/px) was derived from a manually marked reference distance (arena width, 25 cm).

##### Centroid model

U-Net backbone (in_channels=1, kernel_size=3, filters=16, filters_rate=2.0, max_stride=16, stacks=1, convs_per_block=2, output_stride=2); anchor_part=Center, sigma=2.5; total params=1,953,105.

##### Centered-instance model

U-Net backbone (filters=24, filters_rate=2.0, max_stride=16, stacks=1, convs_per_block=2, output_stride=4); anchor_part=Center, sigma=2.5, loss_weight=1.0; crop_size=240; total params=4,311,542. Skeleton: 14 nodes (Nose, Left_ear, Right_ear, Spine_1, Spine_2, Center, Left_fhip, Right_fhip, Left_bhip, Right_bhip, Tail_base, Tail_1, Tail_2, Tail_3).

##### Training (both models)

Adam (lr=0.0001, amsgrad=false), batch_size=4, up to 200 epochs (200 steps/epoch), validation_fraction=0.1; ReduceLROnPlateau (patience=5, factor=0.5, threshold=1e-6, cooldown=3, min_lr=1e-8); early stopping (patience=20, min_delta=1e-8). Preprocessing: grayscale, max_height=400, max_width=440, scale=0.8. Augmentation: rotation ±180° (p=1.0), scale 0.9–1.1 (p=1.0); intensity augmentation and mixup disabled.

##### Inference

tracker=simplemaxtracks; match=Hungarian; similarity=centroid; max_tracks=target_instance_count=2; track_window=45; robust=0.95; min_match_points=1; min_new_track_points=2; post_connect_single_breaks enabled; optical flow window=31, max levels=5.

All videos underwent manual proofreading to correct identity swaps (primarily from occlusion by enrichment items), blind to genotype.

### **b.** DeepOF

Corrected tracking files were processed with DeepOF v0.8.6 (Windows workstation, NVIDIA Tesla A100, 40 GB) using a reduced 11-node skeleton (Tail_1–3 excluded for low tracking accuracy), yielding per-frame supervised behavioral classifications and derived .csv outputs.

Principal component analysis was performed on 13 DeepOF-derived behavioral features (six individual/locomotor behaviors: climb-arena, sniff-arena, immobility, stat-lookaround, stat-active, stat-passive; speed; three mutual behaviors: nose2nose, sidebyside, sidereside; three directional behaviors: following, nose2tail, nose2body), separately for each of three time windows (0–6 h, 7–12 h, 0–12 h). Features (percentage of time per behavior, or mean speed) were standardized (z-score) prior to PCA.

Shannon entropy (bits) was computed per animal from the 12 behavioral time-percentage features (‘speed’ parameter excluded, to avoid confounding by MUT locomotor dysfunction), treated as an unnormalized distribution: values were clipped at zero, normalized to sum to 1, and entropy calculated as −Σp·log₂(p) over non-zero entries, reflecting the evenness of an animal’s behavioral repertoire. Distance to centroid in PCA space (PC1–PC2) — computed on the full 13-feature set, including speed — was calculated per animal relative to its own dyad group’s centroid, separately for each time window.

### Exploratory activity dynamics

For each animal, an ’Exploratory State’ was defined per frame as a binary indicator, set to 1 if the animal was classified by DeepOF as engaging in any of five behawior traits (’climb-arena’, ’sniff-arena’, ’stationary-look-around’, ’stationary-active’, ’sniffing’) in that frame, and 0 otherwise. A complementary ’Nest/Not-detected’ state was defined analogously from frames in which the animal’s pose could not be tracked (DeepOF “missing” output, additionally filtered as ≤9 detected nodes based on visual distinction; typically corresponding to the animal being inside or under nest material). Both states were computed per animal in 120-s bins across the 12-h recording, as the percentage of frames within each bin classified as Exploratory or Nest/Not-detected, respectively.

Per-mouse linear trends in Exploratory State and Nest/Not-detected state (**Exploratory Slope** and **Nest Slope**, %/h) were fitted by ordinary least-squares (OLS) regression (scipy.stats.linregress) on bins with valid data. The **Exploratory Shift** was defined as the time point (h) at which each animal’s individually fitted Exploratory State and Nest/Not-detected trend lines intersected. The **Behavioral Instability Index** was defined as the variance of the residuals between each animal’s binned Exploratory State values and its own individual OLS trend line. For visualization purposes only (Fig. 4A–D), group-level trend lines were additionally plotted by regressing the group-averaged Exploratory State and Nest/Not-detected curves; because these visualization trend lines combine group-averaged slopes with intercepts derived from a separate regression on the averaged curves, their visual crossing point does not necessarily coincide with the group’s reported Exploratory Shift value particularly in groups with higher inter-animal variability.

### Classical behavioral paradigms

#### 3-Chamber Social Interaction Test

Social interaction was evaluated under red light in a three-chamber transparent plexiglass arena (45 × 20 × 25 cm per compartment, 3.5 × 3.5 cm central openings) as described previously (Dalbayler et al., 2025, Moy et al., 2004) with minor adjustments. Outer compartments contained wire-mesh enclosures.

Each session began with a 5-minute habituation phase with free access to the empty arena. The test mouse was then temporarily confined to the central compartment while stimuli were positioned. Once released, the mouse was granted 10 minutes of unimpeded access to all chambers. Time spent in each compartment and interaction zone surrounding the enclosures was quantified to assess social preference.

Testing was performed across three sessions at weekly intervals, evaluating three different conditions:

a. Empty vs. Unknown: A novel C57BL/6J male intruder was randomly assigned to one wire enclosure.
b. Empty vs. Known: A cage-mate was placed in one wire enclosure.
c. Unknown vs. Known: A new, unfamiliar conspecific was placed in one enclosure, and a cage-mate in the other.

### Nest parametrization

#### SAM2 nest material segmentation

Nest material was segmented frame by frame across each 12 h recording (25 fps) using custom software built on SAM2, applied zero-shot without fine-tuning. Frames affected by occlusion or segmentation artefacts were identified by automated quality control and excluded from direct detection, and gaps were reconstructed from adjacent valid detections. For each frame, the software returned nest-material contours, a segmentation confidence score and the mask area. Structural descriptors of the nest material (area, number of fragments, solidity and circularity) were derived from these contours. Solidity reflects how compact the material is relative to its convex hull, circularity how closely its outline approximates a circle and fragmentation depicting number of detected masks. Occlusion of nest material by the animals was determined from SLEAP pose-estimation data. Changes in each descriptor were expressed relative to a per-recording baseline defined at the start of the recording, and peak values (95th percentile) of fragmentation, solidity drop and circularity drop were used for group comparisons.

#### Nest-building detection and kinetics

Nest-building bouts were detected per frame per animal by combining SAM2-derived nest contours, SLEAP pose tracking, and DeepOF-derived behavioral states. Detection was based on the animal’s proximity to the nest material, its behavioral state, and its location outside the feeder and water-dispenser zones. Detection criteria were calibrated against manually verified nest-building episodes. Nest-building time was expressed as the percentage of time spent building within consecutive time bins across the 12-h recording. Per-mouse linear trends (Building_Slope, %/h) were fitted by OLS regression on bins with sufficient valid data (>10 bins). The Building Instability Index was the variance of each mouse’s binned values around its own trend line; a normalized version (√variance / mean building time) was used for group comparisons, as the raw index correlated with overall building level.

#### Deacon’s scale and nest weight

Nest weight (including unbuilt baseline material, 2 g; raw) was measured at the end of the 12h recording. Deacon scale scoring was performed by a single rater, blind to genotype, at the end of the 12h recording immediately following video acquisition.

### Statistics

#### Outlier-corrected comparisons

Outliers were identified independently within each group by the IQR method (values <Q1−1.5×IQR or >Q3+1.5×IQR), excluded from statistical testing, and shown in figures for transparency only. Normality was assessed per group (Shapiro-Wilk) on outlier-corrected data; a one-way ANOVA (Tukey HSD post-hoc if significant) was used when all groups were normally distributed, otherwise Kruskal-Wallis (Dunn’s test with Holm correction).

#### *PERMDISP-based comparisons* (distance-to-centroid, convex-hull area)

No outlier exclusion was applied. Global dispersion differences were assessed by a permutation-based F-statistic (PERMDISP; 5000 permutations, seed=42); pairwise comparisons used a permutation test on the absolute difference in group means or hull area, with Holm correction.

*Paired comparisons* (within-cage nest-building time; SAM2 confidence in occluded vs. non-occluded frames within the same recording): Wilcoxon signed-rank test.

#### Correlation

Pearson correlation; within-genotype correlations computed separately to control for genotype as a confound.

#### *Ordinal data* (Deacon scale)

Kruskal-Wallis on raw scores, followed by Dunn’s test with Holm correction.

#### Segmentation-quality group comparisons

Kruskal-Wallis across the three dyad groups, without outlier exclusion.

Significance throughout: * p<0.05, ** p<0.01, *** p<0.001, **** p<0.0001. Analyses were performed in Python (scipy.stats, statsmodels, scikit-posthocs).

## Results

### Partner-based social behavior dynamic

To characterize the dynamics of autonomous and social behavior mice were recorded continuously for 12h (the full active phase) in a home-cage-like arena (Fig. 1A). Each cage contained two animals in three possible dyad combinations: WT – WT, WT - MUT and MUT - MUT (Fig. 1), rendering four possible assignments of individuals to the following groups: WT (with WT), WT (with MUT), MUT (with WT), MUT (with MUT).

**Fig. 1.**
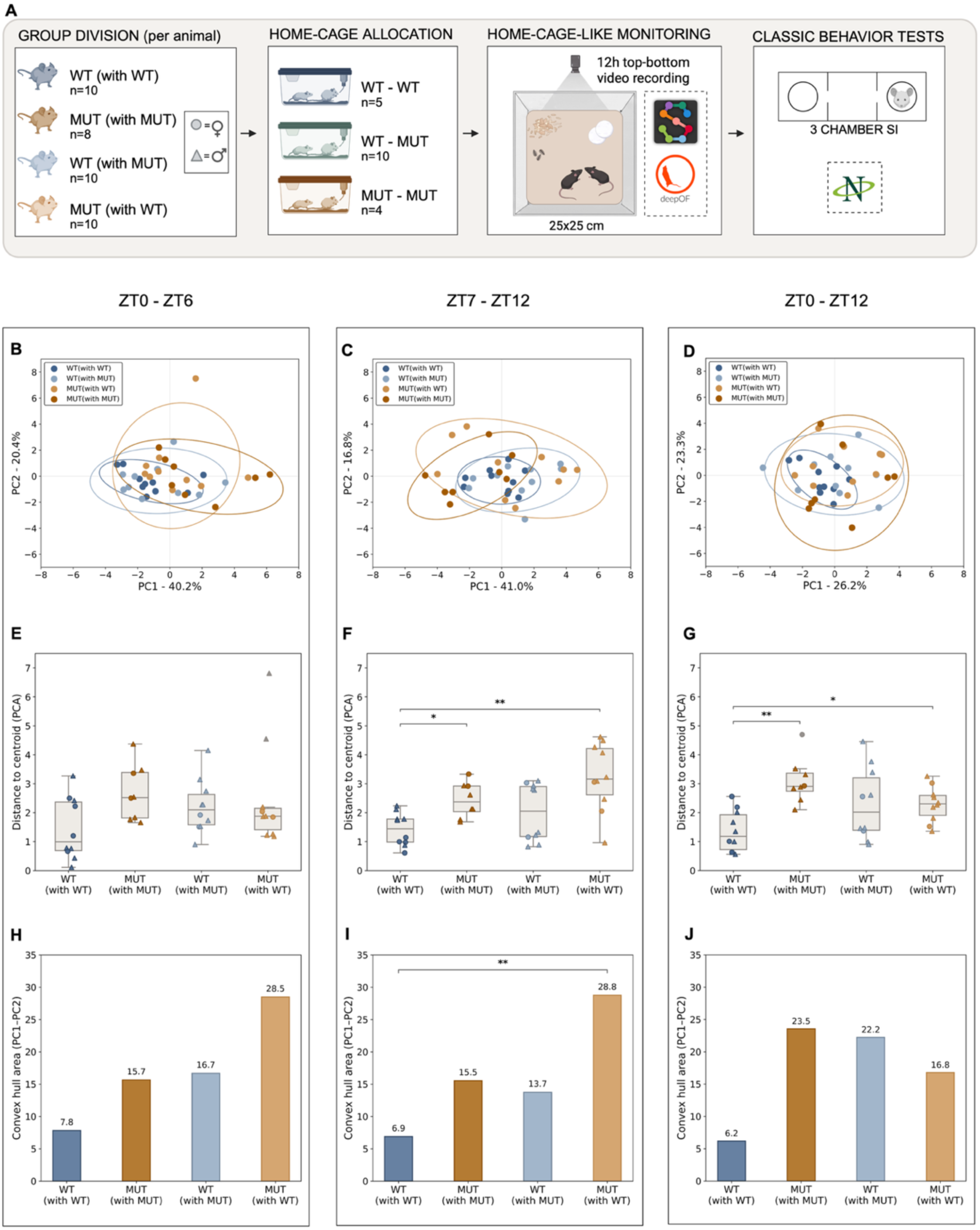
Home-cage-like behavioral profiling reveals time-dependent behavior alterations. **(A)** Experimental design: four dyad groups by individual/partner genotype (WT (with WT), n=10; WT (with MUT), n=10; MUT (with WT), n=10; MUT (with MUT), n=8; cage-level: WT-WT n=5, WT-MUT n=10, MUT-MUT n=4) were housed in pairs and subjected to12-h videorecording of spontaneous behavior, followed by pose estimation (SLEAP) and feature analysis (DeepOF, SAM2). **(B–D)** PCA of DeepOF behavioral features, 0–6 h (B), 7–12 h (C), 0–12 h (D). Ellipses: 95% CI per group. Axes: % variance explained. **(E–G)** Distance to group centroid (PC1–PC2), per animal, samewindows. Boxes: median/IQR; triangles/circles: male/female. PERMDISP (Holm), * p<0.05, ** p<0.01. **(H–J)** Convex hull area (PC1–PC2), same windows. Permutation test (Holm), ** p<0.01; colors: WT (with WT) dark blue; MUT (with MUT) dark orange; WT (with MUT) light blue; MUT (with WT) light orange; graphic created in https://BioRender.com

Behavioral profiles were analyzed with an automated, AI-supported workflow (Jurek et al. 2026), encompassing pose estimation with SLEAP.ai (Pereira et al., 2022), and behavioral interpretation completed with DeepOF (Miranda et al., 2023) and SAM2 (Ravi et al., 2024). Since Shannon entropy has previously been used to quantify the structure and predictability of behavioral repertoires in mice (Shemesh et al., 2013; Peng et al., 2025), we integrated entropy calculated from the relative time spent in each behavioral category as an appropriate proxy for behavioral complexity. Higher entropy reflects a more even, less predictable distribution of time across behaviors, while lower entropy indicates a repertoire dominated by few stereotyped behaviors. Additionally, we tested differences in behavioral heterogeneity using permutational analysis of multivariate dispersion (PERMDISP2). We calculated within-group average distances to the group centroid, where large values indicate high behavioral heterogenity and low values – low behavioral predictability (Carter AJ, Feeney WE, 2012).

Because laboratory mice activity during the active phase is not temporarily uniform (Hasan et al., 2018), behavior was analyzed across three time windows: 0–6 h (ZT0 - ZT6), 7–12 h (ZT7 – ZT12), and 0–12 h (ZT0 – ZT12) to test whether genotype-or partner-based effects were stable or time-dependent across the session.

No strong genotype-or partner-based clustering was observed at the level of overall behavioral profile in any of the three time windows, as assessed with PCA performed on the DeepOF-derived behavioral feature set for each window (Fig. 1B-D). Shannon entropy did not differ between groups in any window (one-way ANOVA; ZT0 – ZT6: p = 0.239; 6–12h: p=0.994; 0–12h: p=0.144; Supp. Fig. 1A–C), indicating that both genotype and dyad composition did not affect the diversity of behaviors displayed by individual subjects. However, within-group dispersion differed significantly between groups in all three windows (ZT0 – ZT6: ANOVA p=0.030; ZT7 – ZT12: Kruskal-Wallis p=0.004; ZT0 – ZT12: ANOVA p=0.007), with WT (with WT) showing the lowest mean dispersion. This analysis indicated that WT (with WT) animals showed the most homogenous, predictable behavioral profile among all four dyad groups. No pairwise comparison reached significance at 0–6h (Tukey HSD, all p-adj>0.05; Fig. 1E–G).

The concentration of dispersion effect in the ZT7 – ZT12 time window, largely absent in ZT0 - ZT6, suggested that a partner-driven behavioral difference was changing over time rather than being present from the start. Independent testing with PERMDISP on the same values confirmed reaching no significance at ZT0 – ZT6 (pseudo-F=1.79, p=0.156) (Carter & Feeney, 2012). To corroborate this pattern, a group dispersion was assessed using convex hull area, a complementary, non-parametric measure of behavioral spread. Both MUT (with WT) and MUT (with MUT) diverged from WT (with WT) specifically in the ZT7 – ZT12 window (MUT (with WT) vs. WT (with WT), p=0.0048; MUT (with MUT) vs. WT (with WT), p=0.017; no other pairs significant. A permutation test on convex hull area identified one significant pair: MUT (with WT) vs. WT (with WT) at 7–12h (p=0.008; Fig. 1H–J).

### Specific behavior traits underlie genotype-and partner-dependent behavioral profile

Having identified partner-driven behavior alterations in the ZT7 - ZT12, mice autonomous behaviors, derived by DeepOF were tested separately for the ZT7 – 12 and ZT0 – ZT12 windows to identify specific underlying traits (Fig. 2A-B).

**Fig 2.**
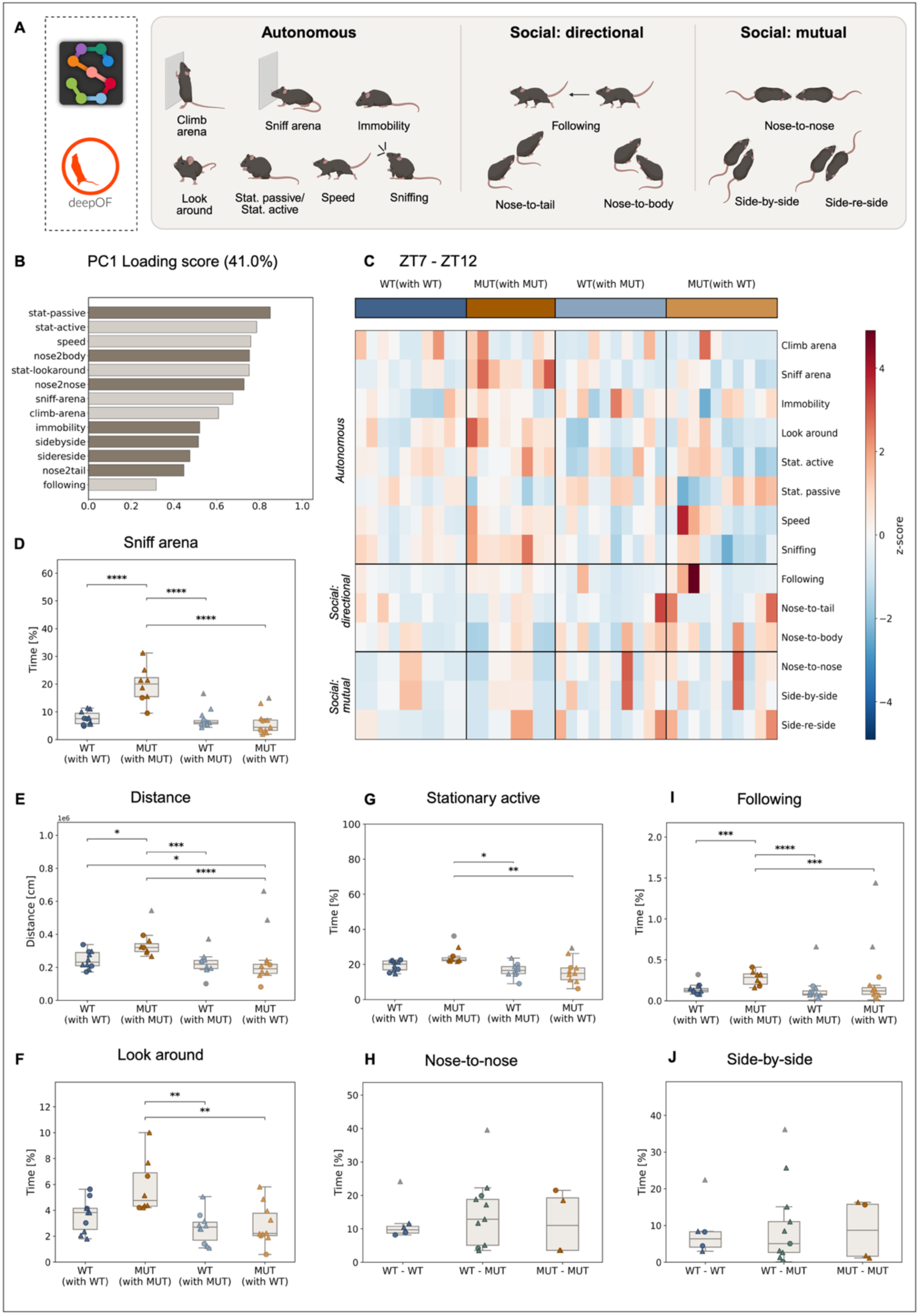
Specific behavior-triats drive behavioral heterogeneity. **(A**) Behavioral traits derived from SLEAP and DeepOF; **(B)** Loading scores of the 13 behaviors on PC1 (41% of variance explained); **(C)** Z-scored per-animal heatmap of the 13 DeepOF-derived behaviors, grouped by category (Autonomous, Social: directional, Social: mutual) and by dyad group (color bar, top); **(D-G)** Boxplots of behaviors differing significantly between groups; **(G–I)** Boxplots of social behaviors showing significance only in ‘Following’ parameter. Boxes show median and interquartile range; triangles and circles depict male and female animals, respectively; gray symbols mark IQR-identified outliers (shown for transparency, excluded from statistical testing, see: Materials and Methods). Global group differences were assessed by one-way ANOVA or Kruskal-Wallis (depending on normality, Shapiro-Wilk); significant pairwise comparisons were determined by Tukey HSD or Dunn’s test with Holm correction, respectively. Asterisks denote post-hoc significance: * p<0.05, ** p<0.01, *** p<0.001, **** p<0.0001; graphic created in https://BioRender.com

This analysis identified parameters of social behavior as being sensitive to the genotype of both, the tested mouse and the partner. PC1 loadings revealed that four behavioral traits differed significantly between groups, with three classified as autonomous behavior: ‘stationary passive’, ‘stationary active’, ‘speed’, and one related to social interaction: ‘nose-to-body’ (Fig. 2C). ‘Sniff arena time’ differed strongly (ANOVA, p<0.0001), with MUT (with MUT) exceeding all three other groups (Tukey HSD, all p<0.0001; Fig. 2D). ‘Total distance moved’ also differed significantly (ANOVA, p<0.0001), with MUT (with MUT) being the most active group (Fig. 2E). Time of looking around’ and ’stationary active’ both differed significantly between groups (’time of looking around’: Kruskal-Wallis, p = 0.003; ’stationary active’: one-way ANOVA, p = 0.016), though driven by distinct patterns — elevated ’time of looking around’ in MUT (with MUT), and reduced ’stationary active’ specifically in MUT (with WT) relative to both WT (with WT) (p = 0.024) and WT (with MUT) (p = 0.032) (Fig. 2F–G). While mutual social behaviors showed no significant differences between groups (Fig. 2G, H, J), the directional social parameter ‘following’ different significantly (p<0.0001), with MUT - MUT pairs showing the highest scores among all dyad compositions (all p≤0.0009; Fig. 2I). This selective effect, confined to a directional rather than mutual social parameter within homecage-like monitoring, motivated further cross-validation using a classical, complementary paradigms.

### Assesing partner-dependent sociabillity in classical 3 Chamber Social Interaction Test

To determine whether partner genotype influences sociability under standardized testing conditions, the 3-chamber social interaction test was performed under three different stages: in the first configuration, WT mice showed significant preference for the compartment with an unknown conspecific (paired t-test, p=0.00024) than the empty one, whereas MUT mice did not show the preference (p=0.473) (Fig. 3A). This strict absence of sociability in MUT stood in contrast to the largely intact social behavior profile observed during long-term monitoring (Fig. 2), suggesting further investigation towards more specific, context-dependent effect.

**Fig. 3.**
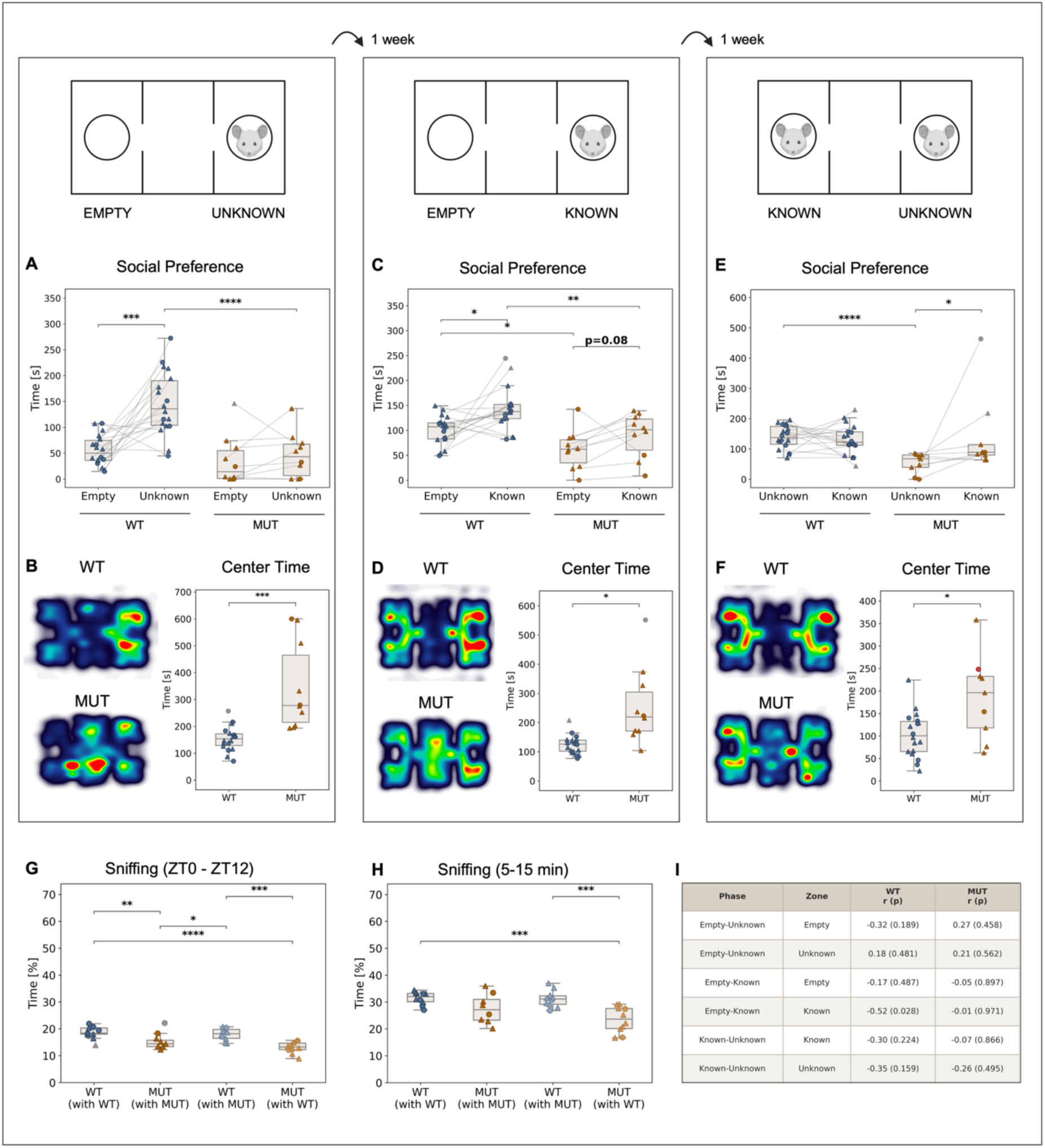
Assessment of sociability in 3 chamber social interaction test. **(A, B)** Empty vs. unknown conspecific (WT n=18, MUT n=10; color code: WT – dark blue, MUT – dark orange, consequently in all pannels). **(A)** Time spent in each zone (Social Preference); **(B)** WT/MUT occupancy heatmaps and time spent in the center zone (Center Time); **(C, D)** Empty vs. known conspecific (cage-mate) WT n=18, MUT n=10; same panel layout as (A, B); **(E, F)** Known vs. unknown conspecific (WT n=18, MUT n=9); same panel layout; **(G, H)** Percentage of time spent sniffing in home-cage-like recordings, across the full 12-h window **(G)** and a 10-min window time-matched to the 3-chamber test (minutes 5–15; **H**), by dyad composition; **(I)** Pearson correlation between sniffing (home-cage-like arena, minutes 5–15) and number of interactions in each 3-chamber zone, computed separately per genotype, N=27–28 (WT: 18, MUT: 9–10); no correlation was considered significant. Heatmaps show mean occupancy density per group (per-animal normalized, Gaussian-smoothed). Boxes: median/IQR; triangles/circles: male/female; gray: IQR-identified outliers (shown, excluded from testing). Paired comparisons (zone vs. zone, within genotype): Wilcoxon signed-rank or paired t-test; between-genotype comparisons: Welch t-test or Mann-Whitney U. * p<0.05, ** p<0.01, *** p<0.001, **** p<0.0001; exact p-value shown where indicated; graphic created in https://BioRender.com

To test whether this reflects unfamiliarity rather than a general social deficit, in the second configuration we used the tested animal’s cage-mate under the wire mesh cup. WT still showed a significant preference for the known conspecific over the empty chamber (p=0.0105), though the effect was weaker than for the unknown conspecific. MUT showed the same directional trend without reaching significance (p=0.084) – an inconclusive result, that warranted a third, more discriminating configuration to determine whether a genuine preference was present.

In the third configuration, WT showed no significant difference in time spent with known versus unknown conspecifics (p=0.625) but approached the unknown conspecific significantly more often (frequency, p=0.0016, Supp. Fig. 3). In contrast, MUT showed a significant preference for the known conspecific (p=0.027), hence WT and MUT differed significantly in time spent with an unknown (p<0.0001), but not with the known conspecific (Fig. 3E). Taken together, results obtained from three configurations indicate that MUT’s apparent lack of sociability (Fig. 3A) does not reflect a general social deficit, but rather a consistent, genotype-driven preference for the familiar over novel conspecific, strengthening as the discriminative contrast between stimuli increases.

We also noticed that MUT mice spent significantly more time in the center compartment of the 3-chamber box than WT. This difference appeared most pronounced in the empty-unknown configuration (p=0.00014) and comparatively smaller once a known conspecific was involved (empty-known, p = 0.01286; known-unknown: Welch t-test, p=0.03162; Fig. 3B, D, F). This data suggests that MUT hesitancy may be particularly enhanced when facing an unfamiliar animal.

To further evaluate possible cause of this reduced social preference, ‘sniffing’ parameter, significantly reduced in MUT in home-cage-like recordings, and determined as genotype-and partner-dependent candidate, was correlated with the ‘number of interactions’ calculated in 3 chamber social tests. This relationship was examined using time-matched window from 12 h recordings (5-15 minutes, mirroring 3-chamber social interaction test structure with 5-min habituation, followed by 10 min of testing phase). No significant correlation was found within either genotype (Fig. 3I). While the small sample size limits statistical power, these results provide no evidence that individual variation in sniffing behavior predicts social approach behavior in either genotype.

### Homecage-like activity dynamics

Having identified autonomous behavioral traits as the primary source of group differentiation (Fig. 2) and given that homecage-like activity dynamics have not previously been characterized in this model, the temporal profile of individual mouse activity was next examined in detail (Fig. 4). This was further motivated by the apparent inconsistency between reduced motor coordination and strength previously reported in R1098Q mice (Miazek et al., 2021; Zalas et al.; 2023) and increased overall distance travelled observed in this study (Fig. 2), suggesting that locomotor features and finer activity dynamics may be dissociable in this model.

**Figure 4.**
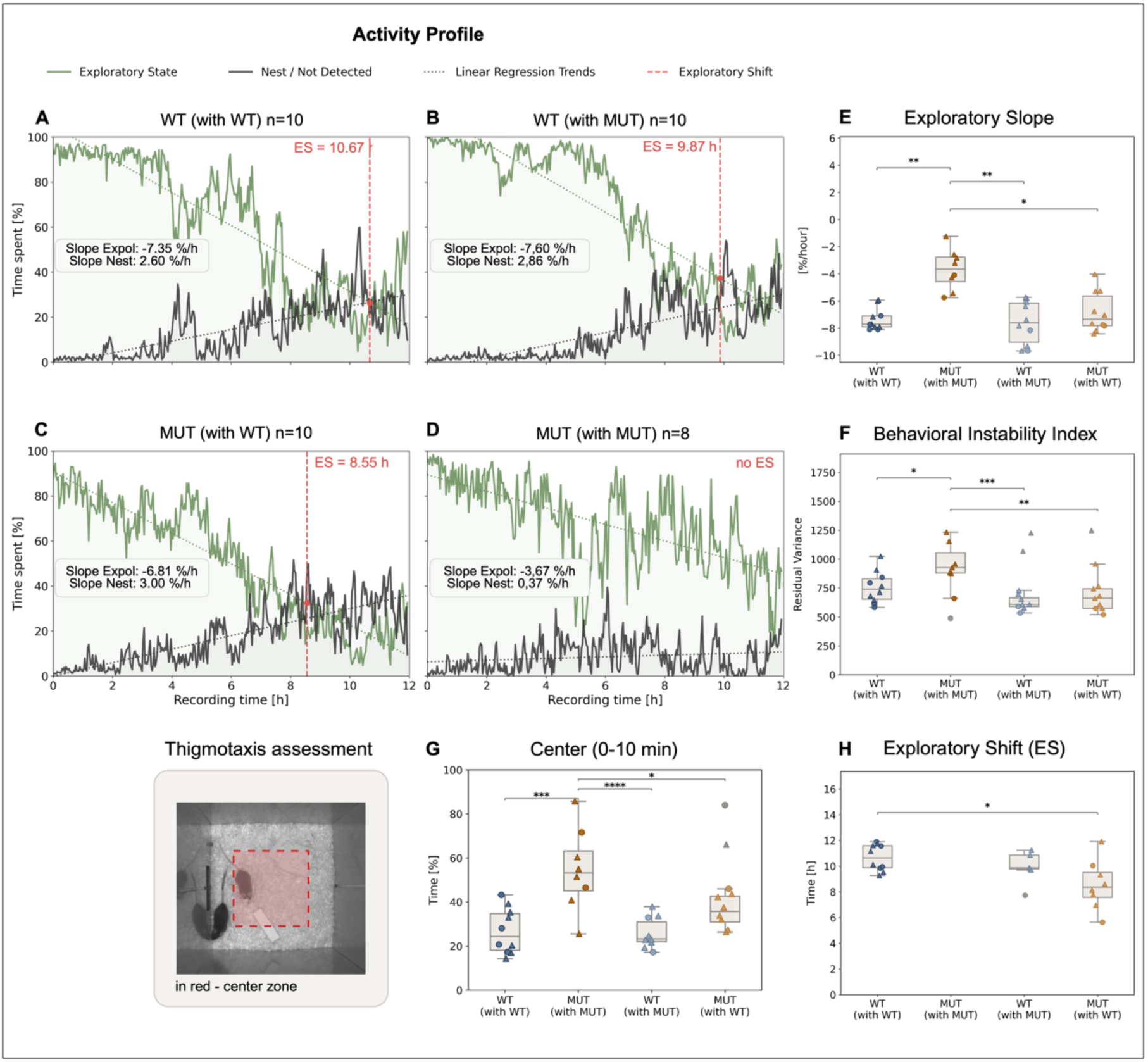
Home-cage activity dynamic. (A–D) Mean exploratory activity profiles (12 h) per mice: **(A)** WT (with WT), **(B)** WT (with MUT), **(C)** MUT (with WT), **(D)** MUT (with MUT); Green: Exploratory State (climb-arena, sniff-arena, stat-look-around, stat-active, sniffing; see Materials and Methods). Black: Nest/not detected. Dotted lines: mean of individual OLS fits; slopes given per panel. Red dashed line: exploratory shift (mean individual Exploratory–Nest intersection); not shown for MUT (with MUT), not reached; **(E)** Exploratory Slope (%/hour); **(F)** Behavioral Instability Index; **(H)** Exploratory Shift timing (h), MUT (with MUT) not shown, not reached; **(G)** Time spent in the center of arena during first 10 minutes of recordings (%). Boxes: median/IQR; triangles/circles: male/female; gray: IQR outliers (shown, excluded from testing). ANOVA or Kruskal-Wallis (normality-dependent); Tukey HSD or Dunn (Holm). * p<0.05, ** p<0.01, *** p<0.001; graphic created in https://BioRender.com

Among the autonomous behaviors examined, a subset specifically describes an animal’s spontaneous exploratory profile, yet each capture only a single, time-avaraged readout of behavior across the full recording. To capture its temporal dynamics instead, a composite ‘Exploratory State’ was derived per animal, and its alterations across full 12 h recordings was summarized using three complementary measures: Exploratory Slope, Behavioral Instability Index and Exploratory Shift (ES) (see Methods; Fig. 4A-D).

Exploratory Slope differed significantly between groups (Kruskal-Wallis, p<0.001): MUT (with MUT) showed the least pronounced decline in exploratory activity over time, differing from all three other groups (vs. WT (with WT): p=0.0023; vs. WT (with MUT): p=0.0013; vs. MUT (with WT): p=0.020), while MUT (with WT) did not differ from either WT (with WT) or WT (with MUT) (both p=1.0; Fig. 4E), suggesting stabilizing effect of the WT partner over MUT.

Consistently, the Behavioral Instability Index (residual variance from each animal’s individual OLS fit) showed MUT (with MUT) differing from all three other groups (vs. WT (with WT): p=0.029; vs. WT (with MUT): p=0.0003; vs. MUT (with WT): p=0.0017). This parameter did not differentiate WT (with MUT) and MUT (with WT) (p=0.868; Fig. 4F), indicating MUT (with MUT) displayed less predictable activity trajectory.

Among the WT (with WT), WT (with MUT) and MUT (with WT) groups, timing of Exploratory Shift (ES) differed significantly overall (ANOVA, p=0.011), driven by a later shift in WT (with WT) relative to MUT (with WT) (p=0.010). Notably MUT (with MUT) was the only group that did not reach the ES within the 12 h window (Fig. 4H).

To further characterize this altered activity profile, the percentage of time spent in the central zone of the arena during the first 10 minutes of recording was additionally calculated, consistent with classical Open Field protocols assessing thigmotaxis. In line with the other indicators described above, MUT (with MUT) spent significantly more time in the center than all other three groups (one-way ANOVA, p < 0.0001; vs. MUT (with WT): p = 0.014; vs. WT (with MUT): p < 0.0001; vs. WT (with WT): p = 0.0001), with other dyads not differing one from another (all p > 0.23) (Fig. G).

Together, these four measures indicate that MUT (with MUT) animals display a more erratic activity profile, with reduced center avoidance, greater instability, and a failure to transition from activity to nest-oriented behavior within the 12 h recording window – an effect largely rescued by the presence of a WT partner. The exception is the Exploratory Shift, which remained significantly earlier in MUT (with WT) relative to WT (with WT) (p = 0.010; Fig. 4). This disrupted activity-to-rest transition motivated a more detailed characterization of nest-building behavior itself, addressed in the following section.

### Development of the new, SAM2-based algorithm for nest material segmentation

Having shown that partner-driven rescue effect did not fully extend to the Exploratory Shift, we next examined whether the presence of a WT partner similarly rescues nest building itself – a behavior previously reported to be impaired in R1098Q mice (Hobbib et al., 2025). Since nest presence alone did not differ significantly between groups (Kruskal-Wallis, p=0.058; see Materials and methods; Supp. Fig. 4), a novel, more sensitive analytical workflow was developed to directly capture nest-building activity (Fig. 5A).

**Figure 5:**
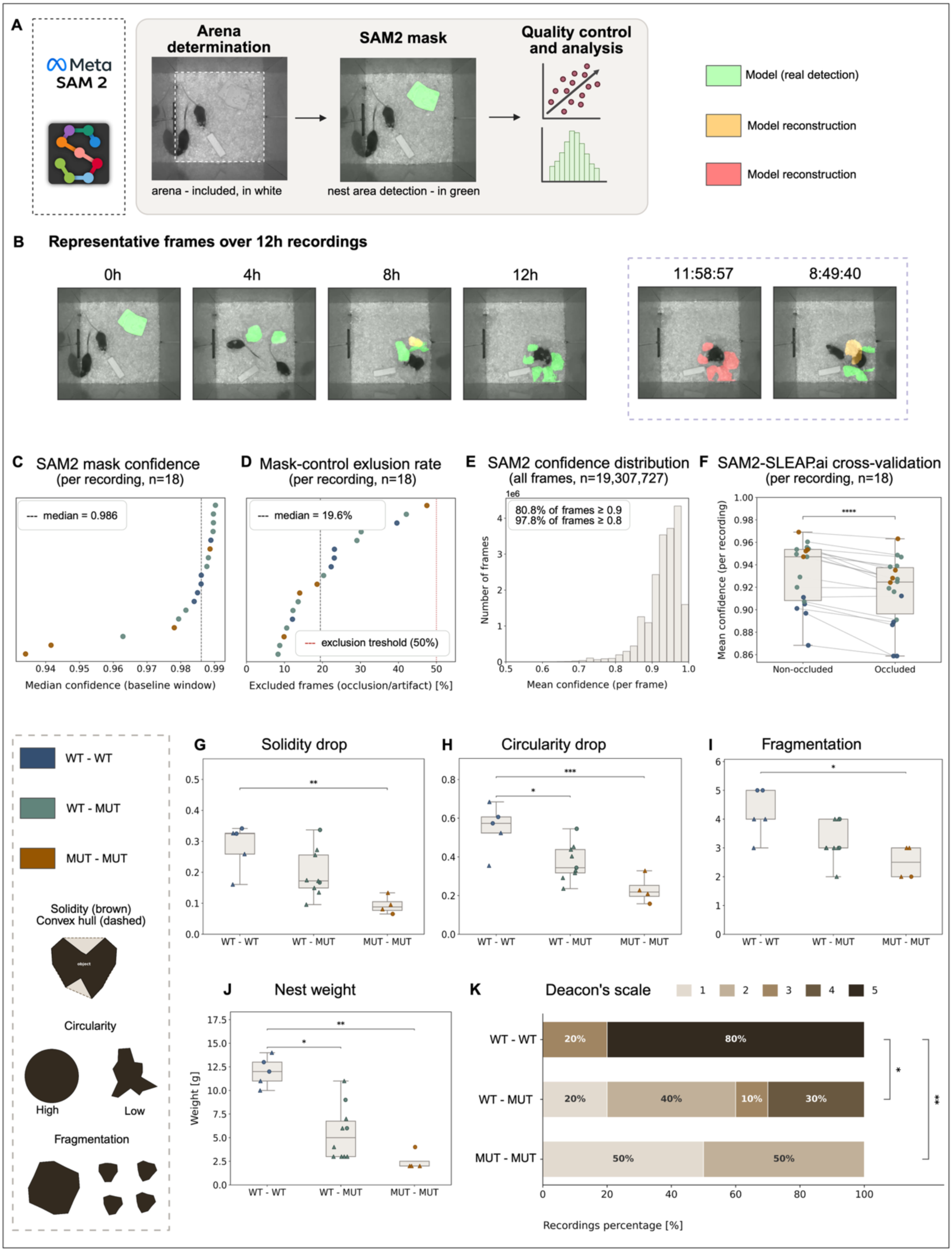
SAM2-based nest material segmentation: pipeline and validation. **(A)** Segmentation pipeline. **(B)** Representative frames, green: model detection; orange and red: model reconstruction. **(C)** Median mask confidence per recording (n=18). **(D)** Excluded-frame percentage per recording (n=18); dotted line: 50% sensitivity threshold. **(E)** Per-frame confidence, all frames (n=19,307,727). **(F)** Confidence in non-occluded vs. occluded frames (SLEAP.ai-defined), paired by recording. **(G)** Peak fragmentation (95th percentile). **(H)** Peak solidity drop (95th percentile). **(I)** Peak circularity drop (95th percentile). Assessed after 12 h recordings: **(J)** Nest weight and **(K)** Deacon scale scoring. Points colored by dyad group. Kruskal-Wallis (C, D, G-K); Wilcoxon signed-rank (F). * p<0.05, ** p<0.01, *** p<0.001, **** p<0.0001; graphic created in https://BioRender.com

Nest material was segmented in each frame of the 12-h recording using custom SAM2-based software (Fig. 5A–B; see Materials and Methods). Segmentation quality was high and consistent across the dataset (n=18 recordings, 19,307,727 frames total). Median per-recording confidence was 0.986 (range: 0.934–0.990; Fig. 5C). Across all frames, 80.8% had a confidence ≥0.90 and 97.8% had a confidence ≥0.80 (Fig. 5E). The proportion of frames excluded from downstream analysis due to occlusion or implausible area changes had a median of 19.5% (range: 8.3–47.5%; Fig. 5D), and no recording exceeded the 50% exclusion threshold used for the sensitivity analysis. Neither confidence nor exclusion rate differed significantly between dyad groups (Kruskal-Wallis; confidence: p=0.13; exclusion rate: p=0.61), indicating that segmentation quality did not depend on dyad composition.

To further validate the segmentation output against an independent data source, per-frame confidence was compared between frames in which the animals occluded the nest material (identified from SLEAP pose-estimation data) and non-occluded frames. Confidence was significantly lower in occluded frames (Wilcoxon signed-rank test, paired by recording; p<0.0001), and non-occluded frames showed higher confidence in 17 of 18 recordings (mean difference: 0.015; Fig. 5F).

Peak fragmentation, solidity drop, and circularity drop – three complementary measures of structural disruption of the nest material – all differed significantly between groups (Kruskal-Wallis; all p <0.05; Fig. 5G-I), consistently driven by greatest material disruption in WT (with WT), relative to MUT (with MUT), with mixed-genotype dyads showing intermediate values. The novel pipeline, introduced in this study, was further cross-validated against nest weight and the Deacon scale, standard approach in nest-quality assessment (Deacon, 2006). Both differed significantly between groups (one-way ANOVA, nest weight: p = 0.0018; Deacon scale: p = 0.0056), following the same pattern as the automatical metrics above: WT – WT outperformed other dyads, while they did not differ from each other (Fig. 5J-K). The convergence between SAM2-based structural metrics with two classical measurements of nest quality provided confirms the validity of automated pipeline against standard nest-quality assessment.

### Nest building kinetics revealed partner-dependent contribution

While nest weight, the Deacon scale and the SAM2-based structural metrics introduced above all capture the endpoint outcome of nest building, none of them alone resolves the dynamic process behind it. To address this gap, nest-building activity was detected for each animal with custom software integrating nest-material segmentation (Fig. 5), SLEAP.ai posture tracking and DeepOF-derived behavioral states (Fig. 6A; see Materials and Methods). Nest-building kinetics were computed per dyad group (Fig. 6B–E), including a Building Instability Index, which quantifies the variability of each animal’s nest-building time around its own individual trend and is analogous to the Behavioral Instability Index introduced previously (Fig. 4).

**Figure 6:**
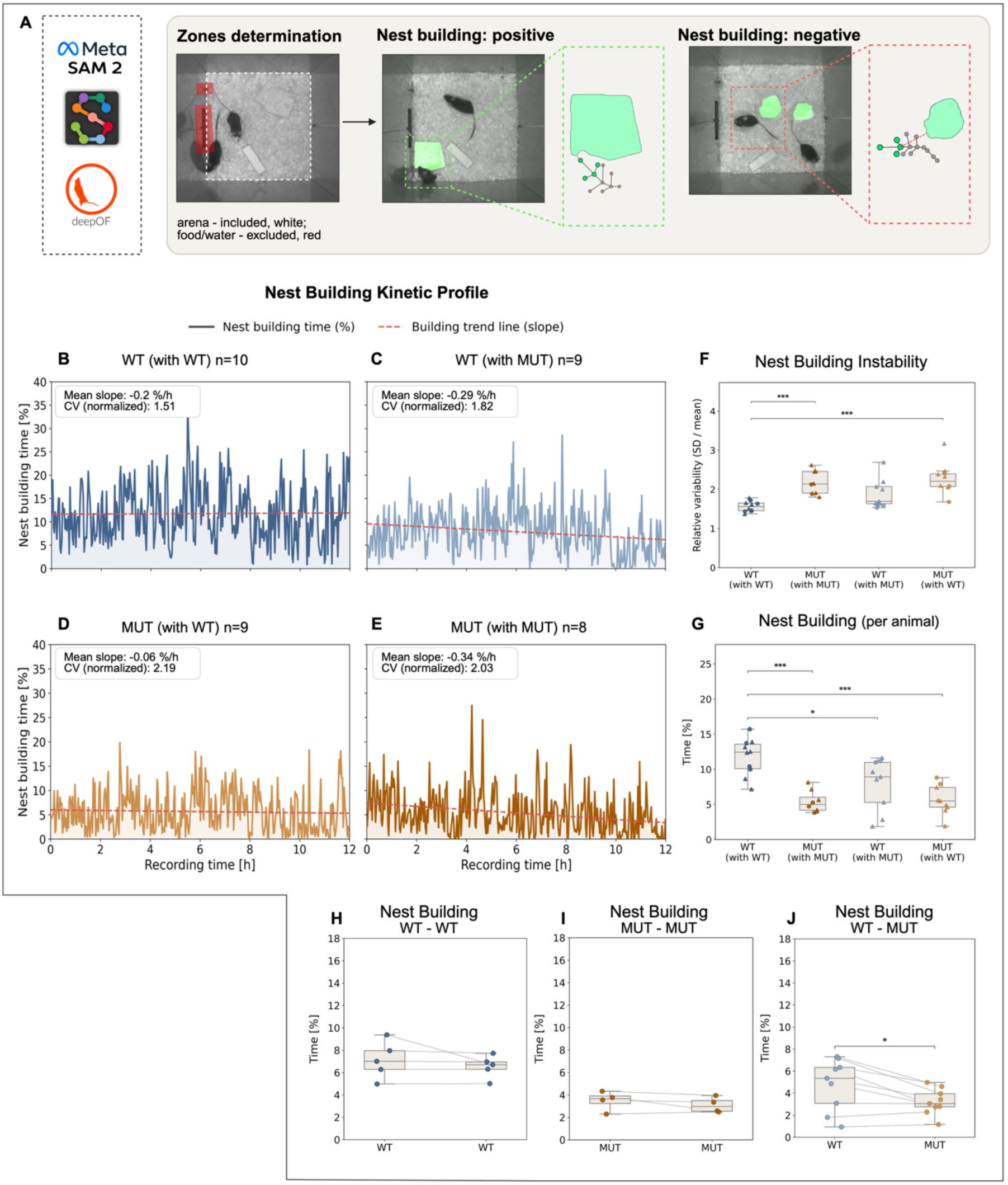
Nest building kinetics revealed partner-dependent contribution. **(A)** Nest-building detection pipeline: zone/prompt definition (SAM2), representative positive and negative frames. **(B–E)** Nest-building time (%) over the 12-h recording per dyad group: WT (with WT) **(B)**, WT (with MUT) **(C)**, MUT (with WT) **(D)**, MUT (with MUT) **(D)**; dotted line: trend (mean slope); CV: normalized coefficient of variation. **(F)** Building Instability (relative variability, SD/mean) across the four dyad groups. **(G)** Nest-building time per animal (4 groups). **(H–J)** Paired nest-building time within the same cage: WT-WT **(H)**, MUT-MUT **(I)**, WT-MUT **(J)**; lines connect cage-mates. Boxes: median/IQR; triangles/circles: male/female; gray: IQR-identified outliers (shown, excluded from testing). Global differences: one-way ANOVA or Kruskal-Wallis (E, I, J); paired Wilcoxon signed-rank (H–J). Post-hoc: Tukey HSD or Dunn (Holm). * p<0.05, ** p<0.01, *** p<0.001; graphic created in https://BioRender.com

Across all four dyad groups, nest-building activity was consistently disrupted by MUT presence in the cage, but in distinct ways. First, Building Instability Index was elevated whenever MUT animal was present in dyad, regardless of partner genotype: MUT (with WT) and MUT (with MUT) pairs showed comparably high instability (p=0.9997), and both differed significantly from WT (with WT) (p=0.0003 and p=0.0004, respectively; one-way ANOVA, p=0.00009; Fig. 6F). While WT (with MUT) occupied an intermediate position, WT (with WT) showed the lowest instability index, indicating, that it is MUT presence that destabilizes the building process itself.

Secondly, within mixed-genotype pairs, the WT animal consistently carried a disproportionate share of building activity. While building time did not differ within cage-mates of the same genotype pairs (paired Wilcoxon signed-rank; WT-WT: p=0.625; MUT-MUT: p=0.625), in mixed-genotype pairs, the WT animal accounted for significantly more building time than its MUT partner (p=0.027; WT higher in 7 of 9 pairs; Fig. 6G–I). This observation was consistent with overall group difference analysis (% of frames, 4 groups; one-way ANOVA, p<0.0001; Fig. 6J). where WT (with WT) differed from all three other groups (vs. WT (with MUT): p=0.016; vs. MUT (with WT): p=0.0001; vs. MUT (with MUT): p=0.0001). Together, this findings indicate that WT animals compensate for their MUT partner’s reduced building contribution, rather than equally contributing to nest formation.

## Discussion

This work reveals previously unknown aspects of central effects of *SPTAN1* R1098Q mouse model, beyond the progressive motor-and memory deficits already characterized in this model (Miazek et al. 2021; Zalas et al., 2023; Hobbib et al., 2025) and points to potential mechanisms of partial rescue of selected deficits through environmental cues. We observed that the phenotype exerted R1098Q mice may be rescued in a domain-dependent manner. The presence of a WT partner in a homecage-like environment rescued selected aspects of a MUT phenotype, such as behavioral dynamics and social interaction, but did not improve cooperative nest-building performance. To place these findings in the broader context of spectrin-related mouse models, Table 1 summarizes behavioral phenotypes reported across mouse models of spectrinopathies, together with the results of the present study.

The impact of cage-mates on autonomous behavior of each individuals with normalization effect on affective-and stress-related domains are well-known. In classical fear-conditioning and acute-stress paradigms, a calm, non-stressed partner reliably buffers conspecific’s behavioral stress response, whereas a stressed partner fails to contage or even exacerbate these responses (Kiyokawa et al., 2004; Beery & Kaufer, 2015). This buffering mechanisms is likely mediated via cingulate-amygdala circuits, homologous to those implicated in rodents’ empathy (Keysers et al., 2022; Gorkiewicz et al., 2022). Notably, such social modulation is not restricted to stress-induced phenotypes: environmental enrichment rescues social deficits in Pten+/-mice, a genetic model of autism spectrum disorder (Clipperton-Allen et al., 2021), and modulates disease severity in Huntington’s disease models (Spires & Hannan, 2005), despite the fully genetic basis of both. Whether a comparable mechanism extends to spontaneous, non-stress-induced dynamics, and which brain circuits might underly it in the *SPTAN1* R1098Q model remains untested. Prior work on this line focused on cerebellar Purkinje cell loss underlying the ataxic phenotype (Miazek et al., 2021) and cortical hyperexcitability in the somatosensory cortex (Hobbib et al., 2025), while limbic circuits implicated in social buffering have not yet been addressed in this mouse model.

In this study, social buffering was first acknowledged at the level of overall behavioral variability: WT (with WT) dyads showed the lowest dispersion (distance to centroid) across all three time windows, indicating that dyad composition shapes behavioral heterogeneity broadly, before specific behavioral traits are considered (Fig. 1). Consistent with this MUT (with MUT) displayed the most erratic and unpredictable activity trajectory of all four groups, while MUT (with WT) did not differ from either WT group (Fig. 4E, F), indicating that a WT partner elevates MUT performance. However, this pattern was not universal: while Fig.4 captured dispersion and instability at the level of ‘Exploratory State’ – a composite of five behavioral traits spanning the full recording, this same buffering effect was not consistently recovered when single behavioral traits were examined individualy within a narrow time window (Fig. 2). That indicates that buffering operates on overall trajectory metrics but does not uniformly extend to each of its underlying components.

Notably, this pattern contrasts with findings in socially isolated mice, where isolation-induced hyperactivity was shown to persist regardless of cohousing with group-reared conspecifics, reflecting a non-social trait unaffected by partner behavior (Endo et al., 2018) — whereas social proximity and approach behavior in that same model were partially rescued by such cohousing. The opposite pattern observed here — activity dynamics being buffered by partner genotype, while a cooperative, socially embedded task (nest-building) was not — suggests that the domain-specificity of social buffering may itself be model-and context-dependent, rather than following a fixed division between "social" and "non-social" behavioral domains.

A reversed buffering pattern was observed in the nest building: rather than WT presence elevating dyad performance, WT - MUT cages were indistinguishable from MUT - MUT cages on endpoint nest-quality measures, differing only from WT-WT (Fig. 6G, H). This data suggests that a MUT partner constrains the WT nest-building performance, rather than being elevated by it. However, nest-building activity is a motor-demanding task, and R1098Q model has previously established deficits in coordination and grip strength (Miazek et al. 2021; Zalas et al., 2023; Hobbib et al., 2025), which themselves may limit MUT contribution independent of any social mechanism. Notably, Hobbib et al., 2025 reported that MUT mice eventually achieve near-normal nest quality (Deacon score 3), following extended, 3-day observation. That assessment, however, involved single-housed animals. Whether the partner-driven constraint on WT contribution observed here would similarly resolve with extended cohousing remains untested.

Those two, non-mutually exclusive explanations may account for this dissociation. First, social-buffering efficacy and direction are known to depend on partner state: calm or behaviorally stable companions reliably buffer distress, whereas distressed or behaviorally atypical conspecifics can fail to buffer, or even exacerbate, negative outcomes in the paired animal (Denommé & Mason, 2022). Under this framework, while a WT partner successfully buffers MUT’s own activity instability, the reverse relationship need not hold as the behaviorally atypical partner, MUT may simply lack the capacity to extend comparable buffering to WT’s joint nest-building output. Second, this deficits could simply reflect a genotype-driven limitation — consistent with previously reported motor coordination, muscle-strength deficits and nest building lower performance in this model (Miazek et al., 2021; Zalas et al., 2023; Hobbib et al., 2025) — largely independent of partner identity. However, a purely genotype-driven account offers no reason for an unaffected WT to underperform, when is paired with MUT. This asymmetry though favors the first explanation, in which the presence of a behaviorally-and genotype-partner constitutes a source of possible social distress that constrains cooperative output, rather than a fixed motor limitation acting independently of partners identity.

Together, these findings indicate that social buffering in this model is domain-and metric-specific rather than globally rescuing, with the underlying genotype-driven dysregulation appearing to be behaviorally modulated, not eliminated by partner state. This is further supported by a convergent, two separate parameters pattern: both distance travelled and following were selectively elevated in MUT (with MUT) dyads and normalized when MUT was paired with WT (Fig. 2). It points to WT-mediated regulatory effect spanning both exploratory-and social approach domain simultaneously. Notably, mutual-and most directional social behavior – with exception of ‘following’ did not differ between dyads or individual animals, suggesting that social behavior is more resilient to *SPTAN1* R1098Q genotype-driven alterations than autonomous locomotor behavior (Fig. 2). The core neural circuitry mediating affiliative social behavior — the social decision-making network, comprising the evolutionarily conserved social behavior network (Newman, 1999) and mesolimbic reward system — has been shown to be conserved across vertebrate taxis (O’Connell & Hofmann, 2012). While this conservation does not by itself predict resistance to any specific genetic perturbation, it is consistent with a general expectation that highly interconnected circuits mediating core affiliative behaviors may be more robust to disruption than circuits supporting individually variable, context-modulated behaviors such as the exploratory activity dynamics observed in our data. Critically, MUT (with MUT) dyads did not avoid or ignore one another; rather, they displayed mutually escalated activity and excessive, unmodulated pursuit, indicating dysregulated rather than diminished social engagement. This is consistent with atypical, poorly modulated social approach reported in some Autism Spectrum Disorder models (ASD), and with self-regulation difficulties (low frustration tolerance, aggression, and impulsivity) reported in a human carrier of a heterozygous *SPTAN1* variant diagnosed with ADHD and autism spectrum disorder, and in ADHD patients carrying autosomal dominant β-II spectrin variants (Luongo-Zink et al., 2022; Zalas et al., 2023). This distinction between intrinsic, genotype-driven dysregulation and its partner-dependent behavioral modulation is relevant to interpreting partner-genotype effects in other spectrinopathy and neurodevelopmental models.

Sociability in the 3 chamber social interaction test has traditionally been inferred from several indirect measures such as time spent in the chamber with unknown/known conspecific (Kumar & Sharma, 2016; Piccin & Contarino, 2020) or number of interactions (sniffing) toward the cage (Juybari et al., 2020; Ferri et al., 2016). Although these variables are often interpreted as indicators of social motivation, they represent distinct behavioural processes, including spatial preference, proximity seeking, and olfactory investigation (Sofyana et al., 2026). WT mice displayed a typical sociability profile, spending more time with both unfamiliar and familiar conspecifics than with an empty chamber, and approaching the unfamiliar conspecific more frequently in the final configuration despite no significant time difference (Fig. 3). This weak cage-mate preference is consistent with reports that inbred C57BL/6J mice show comparatively weak familiar-partner preference even in a fully wild-type context (Dudas et al., 2025). In contrast, MUT mice showed no preference for the unfamiliar conspecific and instead preferred the familiar cage-mate by the final test configuration, displaying a shift in preference strategy (Fig. 3).

This preference for a known conspecific might intuitively be attributed to a social-buffering, calming effect of familiarity on the tested animal; however, familiarity does not uniformly predict physiological stress reactivity or a single behavioral strategy in rodents. A familiar partner more effectively buffers conditioned fear responses than an unfamiliar one (Kiyokawa et al., 2014) and can shift social approach strategies toward or away from a distressed conspecific in a sex-specific manner (Rogers-Carter et al., 2018). This suggests that the behavioral shift in MUT does not rely on impaired capacity for social interaction but may be related to individual emotional regulation or arousal in social contexts. We therefore interpret the observed preference for the known conspecific as reflecting a context-dependent shift in social strategy rather than a straightforward index of reduced stress.

Given that prior work on *SPTAN1* R1098Q model reported selective deficits in novel object recognition (Zalas et al., 2023), reduced sociability might be mistaken for cognitive dysfunction in MUT. However, a comparable dissociation has been reported for another cytoskeletal regulator: Cofilin1-deficient mice. Those animals, which lack a key actin-dynamics regulator, did not show impaired social interaction, social approach, and social recognition alterations, while they displayed selective deficits in novel object recognition, leading the authors to conclude that cofilin1 is relevant specifically for non-social cognition (Sungur et al., 2018). This supports the broader principle that cytoskeletal proteins can be dispensable for core affiliative social behavior while remaining critical for autonomous, non-social cognitive processes — consistent with the pattern we observe for *SPTAN1* R1098Q mice.

MUT mice also spent significantly increased time in the center chamber during all three testing phases, a measure that conflates anxiety, indecision and reduced exploration phenotype (Yang et al., 2011; Conduct Science Three Chamber Sociability Index Calculator, v.1.0) and could in principle indicate anxiety-like profile. However, this is difficult to reconcile with the MUT’s behavioral profile: increased overall distance traveled and the highest exploratory scores of any genotype (Fig. 2) are more consistent with reduced anxiety-like behavior and heightened general activity than with an anxiogenic phenotype. This interpretation directly converges with already described report of the same mouse model -Hobbib et al., 2025 similarly found that MUT mice spend more time in the center of an open field, with reduced thigmotaxis interpreted as a less-anxious phenotype, independently confirmed with our homecage-like settings (Fig. 4G). We therefore favor an interpretation in which the increased center time reflects greater overall locomotion and exploratory drive in MUT mice, rather than avoidance or hesitancy toward either stimulus, consistent with the broader caution that time-based chamber measures conflate spatial preference, proximity-seeking, and motivational state (Sofyana et al., 2026).

Beyond these biological findings, the present homecage-like approach itself offered multiple advantages over classical, single time point testing. Continuous observation of cage-mate dynamics not only indicated the social buffering phenomenon itself, but motivated designing three chamber social interaction test phasing in non-standard configuration. Without subjecting animals to an additional, dedicated open field test to assess thigmotaxy, data could be easily extracted from 12h recordings, giving replicated results obtained in a traditional way (Hobbib et al., 2025). Finally, we introduced a 2D video-based framework for quantitative monitoring of the nest-building process, implemented as custom software. The software achieves reliable structural and temporal resolution without specialized hardware, by cross-referencing the outputs of three independent computational tools: SLEAP for markerless pose estimation of individual animals, DeepOF for supervised classification of autonomous and social behaviors, and SAM2 for frame-by-frame segmentation and tracking of the nest structure itself. While prior work has employed SAM2 as an annotation tool to accelerate the generation of training data for downstream mouse-detection models, and has noted nests specifically as a source of occlusion complicating animal-centric detection (Kaul et al., 2024), to our knowledge no study has used SAM/SAM2 as the deployed, direct segmentation-and-tracking engine for the nest structure itself. By combining animal posture, behavioral state, and nest morphology within a single analytical framework, this approach captures nest-building as dynamic process -including its instability and partner-dependent contribution-rather than static endpoint observation assessed by traditional paradigm. This pipeline provides a scalable, non-invasive alternative for studying nest-building in genetically and socially heterogeneous groups.

## Acknowledgments

P.Z. and A.M. thank the staff of the UPWr Animal Facility, especially Monika Ossowska and Aleksandra Caban for technical support. P.Z. and M.S. are grateful for members of AstroGroup for technical insights and constructive feedback. P.Z is grateful to T. Prószyński for lending the 3 chamber social interaction test setup and to M. Skuła for creative discussions and support. This work was supported with grants obtained from: the Ministry of Science and Higher Education, Poland, „Industrial PhD Programme, VI edition”, number DWD/6/00191/2022 and National Science Centre, Poland, number 2021/41/B/NZ3/04099.

## Authors contribution

P.Z., M.S. and A.M. conceptualized the project and designed the study. P.Z. performed the behavioral experiments, including the three-chamber social interaction test and 12-h homecage-like monitoring, and analyzed the data. P.Z. conceptualized the nest-building parametrization, developed the custom software and performed the analysis of complex behavior. M.S. and A.M. supervised the project. P.Z. wrote the manuscript with input from M.S. and A.M. All authors reviewed and approved the final version of the manuscript.

## Data availability

Data and Supplementary Materials supporting this study are available upon reasonable request

## References

1. Armbrust, K.R., Wang, X., Hathorn, T.J., Cramer, S.W., Chen, G., Zu, T., Kangas, T., Zink, A.N., Oz, G., Ebner, T.J., & Ranum, L.P.W. (2014). Mutant β-III spectrin causes mGluR1 mislocalization and functional deficits in a mouse model of spinocerebellar ataxia type 5. Journal of Neuroscience, 34(30), 9891–9904. 10.1523/JNEUROSCI.0876-14.2014

2. Baines, A.J. (2009). Evolution of spectrin function in cytoskeletal and membrane networks. Biochemical Society Transactions, 37(4), 796–803. 10.1042/BST0370796

3. Beery, A.K., & Kaufer, D. (2015). Stress, social behavior, and resilience: Insights from rodents. Neurobiology of Stress, 1, 116–127. 10.1016/j.ynstr.2014.10.004

4. Carter, A.J., & Feeney, W.E. (2012). Taking a comparative approach: Analysing personality as a multivariate behavioural response across species. PLoS ONE, 7(7), e42440. 10.1371/journal.pone.0042440

5. Clipperton-Allen, A.E., Zhang, A., Cohen, O.S., & Page, D.T. (2021). Environmental Enrichment Rescues Social Behavioral Deficits and Synaptic Abnormalities in Pten Haploinsufficient Mice. Genes, 12(9), 1366. 10.3390/genes12091366

6. ConductScience, Inc. (2026). *Three-Chamber Sociability Index Calculator* (Version 1.0) [Computer software]. https://conductscience.com/tools/three-chamber-sociability-index-calculator

7. Cousin, M.A., Creighton, B.A., Breau, K.A., Spillmann, R.C., Torti, E., Dontu, S., Tripathi, S., Ajit, D., Edwards, R.J., Afriyie, S., Bay, J.C., Harper, K.M., Beltran, A.A., Munoz, L.J., Falcon Rodriguez, L., Stankewich, M.C., Person, R.E., Si, Y., Normand, E.A., … Lorenzo, D.N. (2021). Pathogenic SPTBN1 variants cause an autosomal dominant neurodevelopmental syndrome. Nature Genetics, 53(7), 1006–1021. 10.1038/s41588-021-00886-z

8. Dalbeyler, S., Herud, A., Zglinicki, B., Ziuzia, P., Piechota, M., Hoinkis, D., Bergauer, L., Menacho Pando, C., Soltys, Z., Hanus, P., Groves, P., Dixon, L., Ganglberger, F., Del Prete, D., Dubovyk, V., Aßfalg, M., Violet, C., Lawless, N., Kielbasa, S.M., Sigrist, H., Pryce, C., Turecki, G., Schmidt, M.V., Benes, V., Kuner, T., Korostynski, M., Hengerer, B., & Slezak, M. (2025). Glucocorticoid receptors mediate reprogramming of astrocytes in depression. bioRxiv. 10.1101/2025.04.04.646549

9. Deacon, R.M.J. (2006). Assessing nest building in mice. Nature Protocols, 1(3), 1117–1119. 10.1038/nprot.2006.170

10. Denommé, M.R., & Mason, G.J. (2022). Social Buffering as a Tool for Improving Rodent Welfare. Journal of the American Association for Laboratory Animal Science, 61(1), 5–14. 10.30802/AALAS-JAALAS-21-000006

11. Derbala, M.H., Guo, A.S., Mohler, P.J., & Smith, S.A. (2018). The role of βII spectrin in cardiac health and disease. Life Sciences, 192, 278–285. 10.1016/j.lfs.2017.11.009

12. D’Este, E., Kamin, D., Balzarotti, F., & Hell, S.W. (2016). Subcortical cytoskeleton periodicity throughout the nervous system. Scientific Reports, 6, 22741. 10.1038/srep22741

13. Dudas, A., Novak, A., Gora, C., Pecnard, E., Azzopardi, N., Morisset-Lopez, S., & Pellissier, L.P. (2025). Mixed genetic background better recapitulates developmental and psychiatric phenotypes and heterogeneity than inbred C57BL/6J mice. Scientific Reports, 15(1). 10.1038/s41598-025-26215-6

14. Endo, N., Ujita, W., Fujiwara, M., Miyauchi, H., Mishima, H., Makino, Y., Hashimoto, L., Oyama, H., Makinodan, M., Nishi, M., Tohyama, C., & Kakeyama, M. (2018). Multiple animal positioning system shows that socially-reared mice influence the social proximity of isolation-reared cagemates. Communications Biology, 1, 225. 10.1038/s42003-018-0213-5

15. Ferri, S.L., Kreibich, A.S., Torre, M., Piccoli, C.T., Dow, H., Pallathra, A.A., Li, H., Bilker, W.B., Gur, R.C., Abel, T., & Brodkin, E.S. (2016). Activation of basolateral amygdala in juvenile C57BL/6J mice during social approach behavior. Neuroscience, 335, 184–194. 10.1016/j.neuroscience.2016.08.006

16. Goodman, S.R., Zagon, I.S., & Riederer, B.M. (1987). Spectrin isoforms in mammalian brain. Brain Research Bulletin, 18(6), 787–792. 10.1016/0361-9230(87)90217-6

17. Gorkiewicz, T., Danielewski, K., Andraka, K., Kondrakiewicz, K., Meyza, K., Kaminski, J., & Knapska, E. (2022). Social buffering diminishes fear response but does not equal improved fear extinction. Cerebral Cortex. 10.1093/cercor/bhac395

18. Hasan, S., Foster, R.G., Vyazovskiy, V.V., & Peirson, S.N. (2018). Effects of circadian misalignment on sleep in mice. Scientific Reports, 8, 15343. 10.1038/s41598-018-33480-1

19. He, L., Jia, X., Li, C., Liu, Y., Yu, J., Chen, Z.X., Yang, W., & Liang, X. (2025). 3D behavior phenotyping and multi-object tracking system of mice based on deep learning. Smart Agricultural Technology, 11. 10.1016/j.atech.2025.101022

20. Hobbib, R.L., Vakilzadeh, G., Barriendos, D.F., Del Angel, Y.C., & González, M.I. (2025). Axon initial segment damage and hyperexcitability in a mutant mouse with increased calpain-dependent cleavage of αII-spectrin. Neurobiology of Disease, 217. 10.1016/j.nbd.2025.107156

21. Hong, W., Kennedy, A., Burgos-Artizzu, X.P., Zelikowsky, M., Navonne, S.G., Perona, P., & Anderson, D.J. (2015). Automated measurement of mouse social behaviors using depth sensing, video tracking, and machine learning. PNAS, 112(38), E5351– E5360. 10.1073/pnas.1515982112

22. Huang, C.Y., & Rasband, M.N. (2018). Axon initial segments: structure, function, and disease. Annals of the New York Academy of Sciences, 1420(1), 46–61. 10.1111/nyas.13718

23. Jhuang, H., Garrote, E., Yu, X., Khilnani, V., Poggio, T., Steele, A.D., & Serre, T. (2010). Automated home-cage behavioural phenotyping of mice. Nature Communications, 1(6). 10.1038/ncomms1064

24. Juybari, K.B., Sepehri, G., Meymandi, M.S., Vakili Shahrbabaki, S.S., Moslemizadeh, A., Saeedi, N., Aminzadeh, A., Nozari, M., Khaksari, M., Haghpanah, T., & Bashiri, H. (2020). Sex dependent alterations of resveratrol on social behaviors and nociceptive reactivity in VPA-induced autistic-like model in rats. Neurotoxicology and Teratology, 81, 106905. 10.1016/j.ntt.2020.106905

25. Kaul, G., McDevitt, J., Johnson, J., & Eban-Rothschild, A. (2024). DAMM for the detection and tracking of multiple animals within complex social and environmental settings. Scientific Reports, 14(1). 10.1038/s41598-024-72367-2

26. Keysers, C., Knapska, E., Moita, M.A., & Gazzola, V. (2022). Emotional contagion and prosocial behavior in rodents. Trends in Cognitive Sciences, 26, 688– 706. 10.1016/j.tics.2022.05.005

27. Kiyokawa, Y., Kikusui, T., Takeuchi, Y., & Mori, Y. (2004). Partner’s stress status influences social buffering effects in rats. Behavioral Neuroscience, 118(4), 798– 804. 10.1037/0735-7044.118.4.798

28. Kiyokawa, Y., Honda, A., Takeuchi, Y., & Mori, Y. (2014). A familiar conspecific is more effective than an unfamiliar conspecific for social buffering of conditioned fear responses in male rats. Behavioural Brain Research, 267, 189–193. 10.1016/j.bbr.2014.03.043

29. Kizielewicz, N., Molk, D., Camacho, J., Thuman, D., & Beale, C. (2026). Effects of cage change and nest transfer on artificial intelligence-measured behaviors in male C57BL/6 mice. Journal of the American Association for Laboratory Animal Science, 65(2), 311–323. 10.30802/AALAS-JAALAS-25-157

30. Kojima, M., Otabi, H., Kumanogoh, H., Toyoda, A., Ikawa, M., Okabe, M., & Mizui, T. (2020). Reduction in BDNF from inefficient precursor conversion influences nest building and promotes depressive-like behavior in mice. International Journal of Molecular Sciences, 21(11). 10.3390/ijms21113984

31. Kumar, H., & Sharma, B. (2016). Memantine ameliorates autistic behavior, biochemistry & blood brain barrier impairments in rats. Brain Research Bulletin, 124, 27– 39. 10.1016/j.brainresbull.2016.03.013

32. Lidhar, N.K., Darvish-Ghane, S., Sivaselvachandran, S., Khan, S., Wasif, F., Turner, H., Sivaselvachandran, M., Fournier, N.M., & Martin, L.J. (2021). Prelimbic cortex glucocorticoid receptors regulate the stress-mediated inhibition of pain contagion in male mice. Neuropsychopharmacology, 46(6), 1183–1193. 10.1038/s41386-020-00912-4

33. Lise, S., Clarkson, Y., Perkins, E., Kwasniewska, A., Sadighi Akha, E., Schnekenberg, R.P., Suminaite, D., Hope, J., Baker, I., Gregory, L., Green, A., Allan, C., Lamble, S., Jayawant, S., Quaghebeur, G., Cader, M.Z., Hughes, S., Armstrong, R.J., Kanapin, A., Rimmer, A., Lunter, G., Mathieson, I., Cazier, J.B., Buck, D., Taylor, J.C., Bentley, D., McVean, G., Donnelly, P., Knight, S.J., Jackson, M., Ragoussis, J., & Németh, A.H. (2012). Recessive mutations in SPTBN2 implicate β-III spectrin in both cognitive and motor development. PLoS Genetics, 8(12), e1003074. 10.1371/journal.pgen.1003074

34. Liu, C.-H., & Rasband, M.N. (2019). Axonal spectrins: Nanoscale organization, functional domains and spectrinopathies. Frontiers in Cellular Neuroscience, 13, 234. 10.3389/fncel.2019.00234

35. Liu, C.-H., Seo, R., Ho, T.S.-Y., Stankewich, M., Mohler, P.J., Hund, T.J., Noebels, J.L., & Rasband, M.N. (2020). β spectrin-dependent and domain specific mechanisms for Na+ channel clustering. eLife, 9. 10.7554/eLife.56629

36. Liu, J., Ye, J., Ji, C., Ren, W., He, Y., Xu, F., & Wang, F. (2024). Mapping the behavioral signatures of Shank3b mice in both sexes. Neuroscience Bulletin, 40(9), 1299–1314. 10.1007/s12264-024-01237-8

37. Lorenzo, D.N. (2020). Cargo hold and delivery: Ankyrins, spectrins, and their functional patterning of neurons. Cytoskeleton, 77(5–6), 129–148. 10.1002/cm.21602

38. Lorenzo, D.N., Badea, A., Zhou, R., Mohler, P.J., Zhuang, X., & Bennett, V. (2019). βII-spectrin promotes mouse brain connectivity through stabilizing axonal plasma membranes and enabling axonal organelle transport. PNAS, 116(31), 15686–15695. 10.1073/pnas.1820649116

39. Lorenzo, D.N., Edwards, R.J., & Slavutsky, A.L. (2023). Spectrins: Molecular organizers and targets of neurological disorders. Nature Reviews Neuroscience, 24(4), 195–212. 10.1038/s41583-022-00674-6

40. Luongo-Zink, C., Ammons, C., Al-Ramadhani, R., Logan, R., Ono, K.E., Bhalla, S., Kheder, A., Marcus, D.J., Drane, D.L., & Bearden, D.J. (2022). Longitudinal neurodevelopmental profile of a pediatric patient with de novo SPTAN1, epilepsy and left hippocampal sclerosis. Epilepsy & Behavior Reports, 19, 100550. 10.1016/j.ebr.2022.100550

41. Mededovic, E., Laurentius, V., Wu, Y., Kopaczka, M., Chen, Z., Schulz, M., Tolba, R., & Stegmaier, J. (2025). No free lunch in annotation either: An objective evaluation of foundation models for streamlining annotation in animal tracking. arXiv. http://arxiv.org/abs/2502.03907

42. Miazek, A., Zalas, M., Skrzymowska, J., Bogin, B.A., Grzymajło, K., Goszczynski, T.M., Levine, Z.A., Morrow, J.S., & Stankewich, M.C. (2021). Age-dependent ataxia and neurodegeneration caused by an αII spectrin mutation with impaired regulation of its calpain sensitivity. Scientific Reports, 11(1). 10.1038/s41598-021-86470-1

43. Miranda, L., Bordes, J., Pütz, B., Schmidt, M.V., & Müller-Myhsok, B. (2023). DeepOF: a Python package for supervised and unsupervised pattern recognition in mice motion tracking data. Journal of Open Source Software, 8(86), 5394. 10.21105/joss.05394

44. Mitchell, E.J., Thomson, D.M., Openshaw, R.L., Bristow, G.C., Dawson, N., Pratt, J.A., & Morris, B.J. (2020). Drug-responsive autism phenotypes in the 16p11.2 deletion mouse model: A central role for gene-environment interactions. Scientific Reports, 10(1). 10.1038/s41598-020-69130-8

45. Morrow, J.S., & Stankewich, M.C. (2021). The spread of spectrin in ataxia and neurodegenerative disease. Journal of Experimental Neurology, 2(3). 10.33696/Neurol.2.045

46. Moy, S.S., Nadler, J.J., Perez, A., Barbaro, R.P., Johns, J.M., Magnuson, T.R., Piven, J., & Crawley, J.N. (2004). Sociability and preference for social novelty in five inbred strains: an approach to assess autistic-like behavior in mice. *Genes*, Brain and Behavior, 3(5), 287–302. 10.1111/j.1601-1848.2004.00076.x

47. Netser, S., Haskal, S., Magalnik, H., & Wagner, S. (2017). A novel system for tracking social preference dynamics in mice reveals sex-and strain-specific characteristics. Molecular Autism, 8(1). 10.1186/s13229-017-0169-1

48. Newman, S.W. (1999). The medial extended amygdala in male reproductive behavior: A node in the mammalian social behavior network. Annals of the New York Academy of Sciences, 877, 242–257. 10.1111/j.1749-6632.1999.tb09271.x

49. O’Connell, L.A., & Hofmann, H.A. (2012). Evolution of a vertebrate social decision-making network. Science, 336(6085), 1154–1157. 10.1126/science.1218889

50. Okayama, T., Goto, T., & Toyoda, A. (2015). Assessing nest-building behavior of mice using a 3D depth camera. Journal of Neuroscience Methods, 251, 151–157. 10.1016/j.jneumeth.2015.05.019

51. Otabi, H., Okayama, T., & Toyoda, A. (2020). Assessment of nest building and social interaction behavior in mice exposed to acute social defeat stress using a three-dimensional depth camera. Animal Science Journal, 91(1). 10.1111/asj.13447

52. Peng, J., Yang, D., Xu, C., & Xu, T. (2025). The Neural Basis of Quantity Discrimination in Mice. bioRxiv. 10.1101/2025.07.26.666982

53. Pereira, T.D., Tabris, N., Matsliah, A., Turner, D.M., Li, J., Ravindranath, S., Papadoyannis, E.S., Normand, E., Deutsch, D.S., Wang, Z.Y., McKenzie-Smith, G.C., Mitelut, C.C., Castro, M.D., D’Uva, J., Kislin, M., Sanes, D.H., Kocher, S.D., Wang, S.S., Falkner, A.L., Shaevitz, J.W., & Murthy, M. (2022). SLEAP: A deep learning system for multi-animal pose tracking. Nature Methods, 19(4), 486–495. 10.1038/s41592-022-01426-1

54. Perkins, E.M., Clarkson, Y.L., Sabatier, N., Longhurst, D.M., Millward, C.P., Jack, J., Toraiwa, J., Watanabe, M., Rothstein, J.D., Lyndon, A.R., Wyllie, D.J.A., Dutia, M.B., & Jackson, M. (2010). Loss of β-III spectrin leads to Purkinje cell dysfunction recapitulating the behavior and neuropathology of spinocerebellar ataxia type 5 in humans. The Journal of Neuroscience, 30(14), 4857–4867. 10.1523/JNEUROSCI.6065-09.2010

55. Piccin, A., & Contarino, A. (2020). Sex-linked roles of the CRF1 and the CRF2 receptor in social behavior. Journal of Neuroscience Research, 98(8), 1561–1574. 10.1002/jnr.24629

56. Provenzano, G., Chelini, G., & Bozzi, Y. (2017). Genetic control of social behavior: Lessons from mutant mice. Behavioural Brain Research, 325, 237–250. 10.1016/j.bbr.2016.11.005

57. Puścian, A., Łeski, S., Górkiewicz, T., Meyza, K., Lipp, H.P., & Knapska, E. (2014). A novel automated behavioral test battery assessing cognitive rigidity in two genetic mouse models of autism. Frontiers in Behavioral Neuroscience, 8, 140. 10.3389/fnbeh.2014.00140

58. Ravi, N., Gabeur, V., Hu, Y.-T., Hu, R., Ryali, C., Ma, T., Khedr, H., Rädle, R., Rolland, C., Gustafson, L., Mintun, E., Pan, J., Alwala, K.V., Carion, N., Wu, C.-Y., Girshick, R., Dollár, P., & Feichtenhofer, C. (2024). SAM 2: Segment Anything in Images and Videos. arXiv:2408.00714.

59. Rogers-Carter, M.M., Djerdjaj, A., Culp, A.R., Elbaz, J.A., & Christianson, J.P. (2018). Familiarity modulates social approach toward stressed conspecifics in female rats. PLoS ONE, 13(10). 10.1371/journal.pone.0200971

60. Rusu, A., Chevalier, C., de Chaumont, F., Nalesso, V., Brault, V., Hérault, Y., & Ey, E. (2023). Day-to-day spontaneous social behaviours is quantitatively and qualitatively affected in a 16p11.2 deletion mouse model. Frontiers in Behavioral Neuroscience, 17, 1294558. 10.3389/fnbeh.2023.1294558

61. Saitsu, H., Tohyama, J., Kumada, T., Egawa, K., Hamada, K., Okamoto, N., Konishi, Y., Yamasaki, M., Nishiyama, K., Murakami, Y., Wada, K., Takahashi, S., Fujita, T., Sasaki, M., Manabe, A., Kondo, N., & Matsumoto, N. (2010). Dominant-negative mutations in alpha-II spectrin cause West syndrome with severe cerebral hypomyelination, spastic quadriplegia, and developmental delay. American Journal of Human Genetics, 86, 881– 891. 10.1016/j.ajhg.2010.04.013

62. Shemesh, Y., & Chen, A. (2023). A paradigm shift in translational psychiatry through rodent neuroethology. Molecular Psychiatry, 28(3), 993–1003. 10.1038/s41380-022-01913-z

63. Shemesh, Y., Sztainberg, Y., Forkosh, O., Shlapobersky, T., Chen, A., & Schneidman, E. (2013). High-order social interactions in groups of mice. eLife, 2, e00759. 10.7554/eLife.00759

64. Sofyana, M., Piggins, H.D., Jackson, M.G., & Robinson, E.S.J. (2026). Beyond the barrier: An assessment of social behaviour in mice using a modified 3-chamber test. Physiology & Behavior, 315, 115381. 10.1016/j.physbeh.2026.115381

65. Spires, T.L., & Hannan, A.J. (2005). Nature, nurture and neurology: gene–environment interactions in neurodegenerative disease. The FEBS Journal, 272(10), 2347– 2361. 10.1111/j.1742-4658.2005.04677.x

66. Stankewich, M.C., Gwynn, B., Ardito, T., Ji, L., Kim, J., Robledo, R.F., Lux, S.E., Peters, L.L., & Morrow, J.S. (2010). Targeted deletion of βIII spectrin impairs synaptogenesis and generates ataxic and seizure phenotypes. PNAS, 107(13), 6022–6027. 10.1073/pnas.1001522107

67. Sungur, A.Ö., Stemmler, L., Wöhr, M., & Rust, M.B. (2018). Impaired object recognition but normal social behavior and ultrasonic communication in cofilin1 mutant mice. Frontiers in Behavioral Neuroscience, 12, 25. 10.3389/fnbeh.2018.00025

68. Wang, C.C., Ortiz-Gonzalez, X.R., Yum, S.W., Gill, S.M., White, A., Kelter, E., Seaver, L.H., Lee, S., Wiley, G., Gaffney, P.M., Wierenga, K.J., & Rasband, M.N. (2018). Critical roles of αII spectrin in brain development and epileptic encephalopathy. Journal of Clinical Investigation, 128(6), 2464–2477. 10.1172/JCI95743

69. Wang, X., Bey, A.L., Katz, B.M., Badea, A., Kim, N., David, L.K., Duffney, L.J., Kumar, S., Mague, S.D., Hulbert, S.W., Dutta, N., Hayrapetyan, V., Yu, C., Gaidis, E., Zhao, S., Ding, J.D., Xu, Q., Chung, L., Rodriguiz, R.M., … Jiang, Y.H. (2016). Altered mGluR5-Homer scaffolds and corticostriatal connectivity in a Shank3 complete knockout model of autism. Nature Communications, 7, 11459. 10.1038/ncomms11459

70. Xu, K., Zhong, G., & Zhuang, X. (2013). Actin, spectrin, and associated proteins form a periodic cytoskeletal structure in axons. Science, 339(6118), 452–456. 10.1126/science.1232251

71. Yang, M., Silverman, J.L., & Crawley, J.N. (2011). Automated three-chambered social approach task for mice. *Current Protocols in Neuroscience*, Chapter 8, Unit 8.26. 10.1002/0471142301.ns0826s56

72. Yang, M., Bozdagi, O., Scattoni, M.L., Wöhr, M., Roullet, F.I., Katz, A.M., Abrams, D.N., Kalikhman, D., Simon, H., Woldeyohannes, L., Zhang, J.Y., Harris, M.J., Saxena, R., Silverman, J.L., Buxbaum, J.D., & Crawley, J.N. (2012). Reduced excitatory neurotransmission and mild autism-relevant phenotypes in adolescent Shank3 null mutant mice. Journal of Neuroscience, 32(19), 6525– 6541. 10.1523/JNEUROSCI.6107-11.2012

73. Yang, Y., Lacas-Gervais, S., Morest, D.K., Solimena, M., & Rasband, M.N. (2004). βIV spectrins are essential for membrane stability and the molecular organization of nodes of Ranvier. The Journal of Neuroscience, 24(33), 7230–7240. 10.1523/JNEUROSCI.2125-04.2004

74. Zalas, M., Skrzymowska, J., Miążek, A., & Miazek, A. (2023). Progressive ataxia, memory impairments, and seizure episodes in Spna2 R1098Q mouse variant affecting alpha II spectrin’s scaffold stability. Brain Sciences, 13(2), 261. 10.3390/brainsci13020261

